# Structural basis for Parkinson’s Disease-linked LRRK2’s binding to microtubules

**DOI:** 10.1101/2022.01.21.477284

**Authors:** David M. Snead, Mariusz Matyszewski, Andrea M. Dickey, Yu Xuan Lin, Andres E. Leschziner, Samara L. Reck-Peterson

## Abstract

Leucine Rich Repeat Kinase 2 (*LRRK2*) is one of the most commonly mutated genes in familial Parkinson’s Disease (PD). Under some circumstances, LRRK2 co-localizes with microtubules in cells, an association enhanced by PD mutations. We report a cryo-electron microscopy structure of the catalytic half of LRRK2, containing its kinase, which is in a closed conformation, and GTPase domains, bound to microtubules. We also report a structure of the catalytic half of LRRK1, which is closely related to LRRK2, but is not linked to PD. LRRK1’s structure is similar to LRRK2, but LRRK1 does not interact with microtubules. Guided by these structures, we identify amino acids in LRRK2’s GTPase domain that mediate microtubule binding; mutating them disrupts microtubule binding in vitro and in cells, without affecting LRRK2’s kinase activity. Our results have implications for the design of therapeutic LRRK2 kinase inhibitors.

## Introduction

Parkinson’s Disease (PD) is the second most common neurodegenerative disease, affecting over 10 million people worldwide. Autosomal dominant missense mutations in Leucine Rich Repeat Kinase 2 (*LRRK2*) are a major cause of familial PD^1–4^, and mutations in *LRRK2* are also linked to sporadic cases of PD^5,6^. All PD-linked mutations in LRRK2 increase its kinase activity^7–10^ and increased LRRK2 kinase activity in the context of a wild-type protein is also associated with sporadic cases^11^. LRRK2-specific kinase inhibitors have been developed to treat PD and are in clinical trials (clinicaltrials.gov).

While it remains unclear how LRRK2 drives PD, LRRK2 has been functionally linked to membrane trafficking^12–14^. Mutant LRRK2 causes defects in endo/lysosomal, autophagosomal, and mitochondrial trafficking^15–19^, and LRRK2 regulates lysosomal morphology^20–23^. Although the bulk of LRRK2 is found in the cytosol, it can also associate with membranes under some conditions^20,21,24–26^. A subset of Rab GTPases, which are master regulators of membrane trafficking^27^, are phosphorylated by LRRK2, and PD-linked LRRK2 mutations increase Rab phosphorylation in cells^9,28^. Phosphorylation of Rabs by LRRK2 is linked to alterations in ciliogenesis^25–27^ and defects in endolysosomal trafficking^14^. LRRK2 also co-localizes with microtubules in cells and in vitro^29–32^. Cellular localization of LRRK2 with microtubules is seen with elevated expression levels and this is enhanced by Type-1 LRRK2-specific kinase inhibitors^29–31,33,34^. In vitro, the catalytic half of LRRK2 alone can bind to microtubules^31^. In addition, many PD-linked mutations (R1441C, R1441G, Y1699C, and I2020T) increase microtubule association in cells^29,33^. It is currently not understood how LRRK2 perturbs cellular trafficking or how the cellular localization of LRRK2—cytosolic, membrane-associated, and/or microtubule-bound—contributes to its function and to PD pathology. Developing tools that control the localization of LRRK2 in cells will be crucial for determining LRRK2’s cellular function and for understanding the molecular basis of PD.

To develop such tools, structural information about LRRK2 is essential. LRRK2 is a large, multidomain protein (Fig. 1a). The amino-terminal half contains armadillo, ankyrin, and leucine-rich repeat domains. The carboxy-terminal half contains LRRK2’s enzymatic domains—both a Roco family GTPase (Ras-Of-Complex, or ROC domain) and a kinase—as well as a scaffolding domain (C-terminal Of Roc, or COR) and a WD40 protein interaction domain. The COR domain is further subdivided into COR-A and COR-B moieties. Here we refer to the catalytic half of LRRK2 as LRRK2^RCKW^, named for its <u>R</u>OC, <u>C</u>OR, <u>k</u>inase and <u>W</u>D40 domains. Recent structures of LRRK2 have revealed the architecture of LRRK2 at near-atomic resolution^31,35^. A 3.5Å structure of LRRK2^RCKW^ showed that LRRK2’s catalytic half is J-shaped, placing the kinase and GTPase domains in close proximity^31^. Later, a 3.5Å structure of full-length LRRK2 revealed that the N-terminal half of LRRK2 wraps around its enzymatic half, with the leucine rich repeats blocking the kinase’s active site in what appears to be an autoinhibited state^35^. In addition to these structures of soluble LRRK2, a 14Å structure of LRRK2 carrying the I2020T PD mutation bound to microtubules in cells was obtained using cryo-electron tomography (cryo-ET)^30^. The cryo-ET map was used to guide integrative modeling, leading to a molecular model for the enzymatic half of LRRK2 bound to microtubules^30^. This model was refined when the 3.5Å cryo-EM structure of LRRK2’s catalytic half was docked into the cryo-ET structure^31^. In these models the C-terminal half of LRRK2 (LRRK2^RCKW^) wraps around the microtubule using two dimerization interfaces, one between WD40 domains and the other between COR-B domains^31^. In addition, in the models the ROC GTPase domain faces the microtubule, although the cryo-ET structure did not reveal any direct interactions between LRRK2 and the microtubule^30^. An isolated ROC domain has also been shown to interact with alpha and beta-tubulin heterodimers^36^.

**Figure 1.**
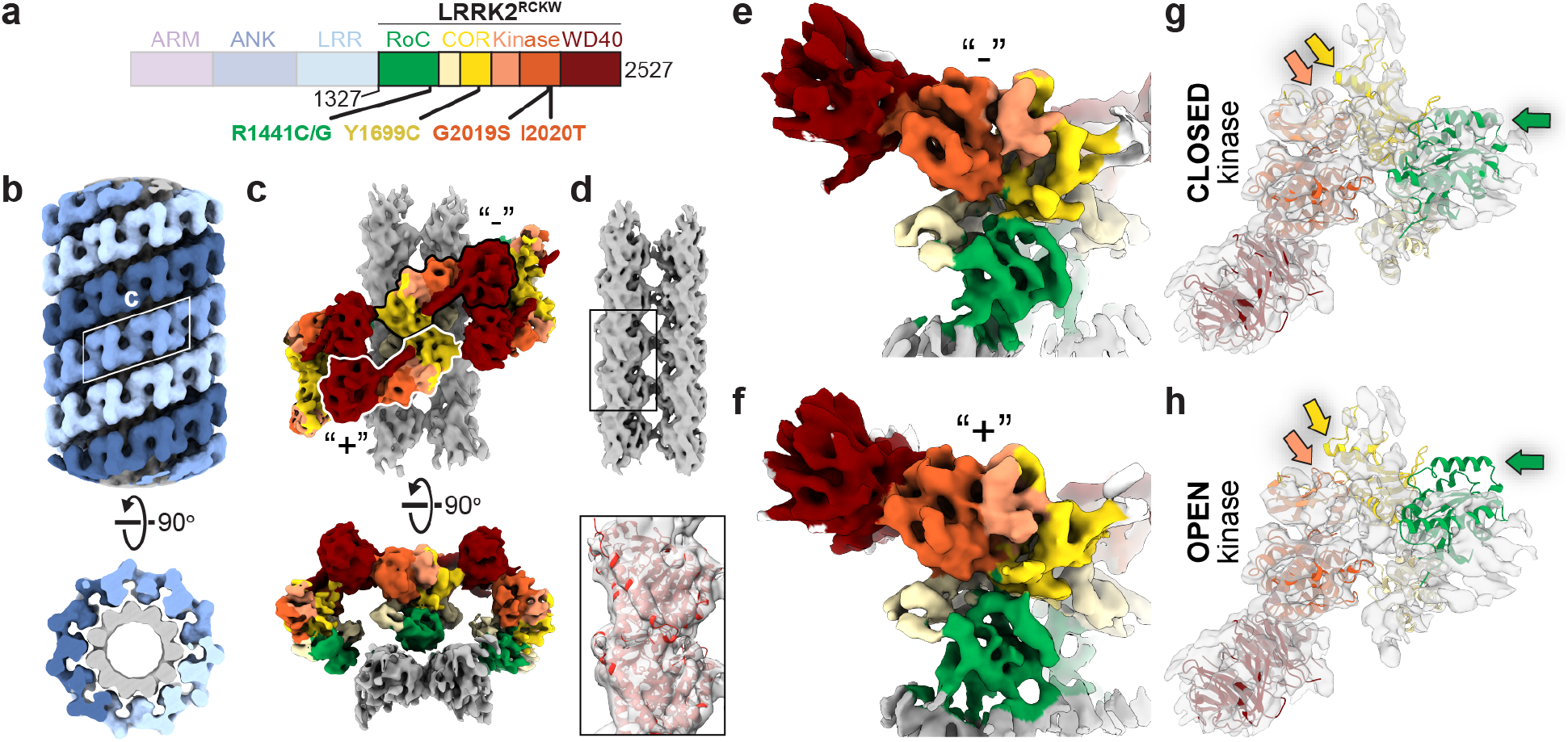
Cryo-EM structure of microtubule-associated LRRK2^RCKW^[I2020T]. **a**, Primary structure of LRRK2. The N-terminal half of LRRK2, absent from the construct used in our cryo-EM studies, is shown in dim colors. The same color-coding of domains is used throughout the Article. **b,** Helical reconstruction (18Å) of LRRK2^RCKW^ [I2020T] filaments bound to a microtubule in the presence of MLi-2. The three LRRK2^RCKW^[I2020T] helices are indicated in different shades of blue. **c**, Cryo-EM reconstruction (6.6Å) of a LRRK2^RCKW^ tetramer and associated microtubule (2 protofilaments), as indicated by the white rhomboid in (b). **d**, Focused refinement of the microtubule in (c) to improve its resolution and determine its polarity. An α/β tubulin dimer (from PDB: 6O2R) was docked into the density (black rectangle and inset below). **e,f**, Focused refinement of the “−” (5.2Å) and “+” (5.0Å) LRRK2^RCKW^ [I2020T] monomers (as labeled in (c)). **g**, The LRRK2^RCKW^ domains (ROC, COR-A, COR-B, Kinase N-lobe, Kinase C-lobe, WD40) (PDB:6VNO) were fitted individually into the 4.5 Å cryo-EM map. **h**, The full LRRK2^RCKW^ model (PDB:6VNO) was aligned to the C-lobe of the kinase as docked in (g). The colored arrows in (g) and (h) point to parts of the model (PDB:6VNO) that fit into the cryo-EM density when domains are docked individually, allowing the kinase to be closed, (g) but protrude from it when the full model is used, which has its kinase in an open conformation (h).

To investigate the possible functional consequences of LRRK2’s interaction with microtubules, we previously looked at the impact of LRRK2 on the movement of microtubule-based motor proteins in vitro^31^. Dynein and kinesin motors move on microtubules, with dynein moving in one direction (towards the microtubule minus end) and kinesin generally moving in the opposite direction (towards the microtubule plus end). Both dynein and kinesin motors interact with their membranous cargos directly or indirectly via connections to Rab GTPases, including those Rabs phosphorylated by LRRK2^37–40^. Using single-molecule assays, we showed that low nanomolar concentrations of LRRK2^RCKW^ blocked the movement of both dynein and kinesin on microtubules^31^. Furthermore, we showed that the conformation of LRRK2’s kinase domain was critical for this effect^31^. LRRK2 predicted to have its kinase domain “closed” (the canonical active conformation) by LRRK2-specific Type-1 kinase inhibitors blocked motility^31^, in agreement with studies showing that these inhibitors enhance the association of LRRK2 with microtubules in cells^29–31,33,34^. In contrast, LRRK2 no longer robustly blocked the movement of dynein or kinesin when its kinase domain was predicted to be in an “open” or inactive conformation (in the presence of Type-2 kinase inhibitors)^31^.

Despite these insights, the structural basis for the formation of LRRK2 filaments and how they interact with microtubules remains unknown. Here, we report a range of cryo-EM structures capturing different levels of detail of the microtubule-bound filaments formed by the C-terminal half of LRRK2 (LRRK2^RCKW^). By focusing our refinements, we were able to resolve the kinase at 4.5Å resolution, the interactions between the ROC domain and the microtubule at 5.0Å, and a LRRK2^RCKW^ tetramer revealing dimerization interfaces at 5.9Å. Our structure reveals direct interactions between LRRK2’s ROC domain and the microtubule. We show that microtubule binding is mediated by electrostatic interactions and requires the negatively charged, glutamate rich carboxy-terminal tubulin tails. We also present a 5.8Å map of the carboxy-terminal half of LRRK1 (LRRK1^RCKW^), LRRK2’s closest human homolog. Despite the structural similarity to LRRK2^RCKW^, we show that LRRK1^RCKW^ does not bind to microtubules. Based on our structure of microtubule-bound LRRK2^RCKW^ and a comparison of our LRRK2^RCKW^ and LRRK1^RCKW^ structures, we identify microtubule-facing basic amino acids that are only conserved in LRRK2’s ROC domain and are required for LRRK2’s interaction with microtubules in vitro and in cells. Mutation of these amino acids reduces LRRK2’s ability to block the movement of kinesin motors in vitro. Together, our work reveals the structural basis for LRRK2’s ability to form filaments and interact with microtubules and identifies mutations that perturb LRRK2’s ability to form filaments and localize to microtubules in cells. These are essential tools for determining the cellular functions of LRRK2 and for the further development of therapeutic LRRK2 kinase inhibitors.

## Results

### Cryo-EM structure of microtubule-associated LRRK2^RCKW^

To understand, structurally and mechanistically, how LRRK2 oligomerizes around and interacts with microtubules, we set out to obtain a higher resolution structure of microtubule-associated LRRK2 filaments using an in vitro reconstituted system and single-particle cryo-EM approaches. We chose to work with LRRK2^RCKW^ because it is sufficient to from filaments in vitro^31^ and it accounted for the density observed in the cryo-ET reconstruction of full-length LRRK2 filaments in cells^30^.

As previously observed^31^, copolymerization of tubulin with LRRK2^RCKW^—either wild type (WT), or carrying the PD-linked mutations G2019S or I2020T—yielded microtubules decorated with LRRK2^RCKW^ (Extended Data Fig.1a). Diffraction patterns calculated from images of these filaments showed layer lines indicating the presence of ordered filaments (Extended Data Fig.1a). In the presence of MLi-2 we saw an additional layer line of lower frequency for all three constructs, suggesting that the filaments had longer-range order (Extended Data Fig.1a). Unlike WT and G2019S, I2020T showed this additional layer line in the absence of MLi-2 as well (Extended Data Fig.1a). Given these observations, we chose to focus on the LRRK2^RCKW^ [I2020T] filaments formed in the presence of MLi-2 for our cryo-EM work. The symmetry mismatch between microtubules, which are polar left-handed helices, and the LRRK2 filaments, which are right-handed and have pseudo-two-fold axes of symmetry perpendicular to the microtubule, required that we largely uncouple their processing (Extended Data Fig.1b,c and Methods). Our cryo-EM analysis resulted in several maps originating from an initial reconstruction of the filaments (Fig.1b): a map of a LRRK2^RCKW^ tetramer that includes density for two microtubule protofilaments (6.6Å) (Fig.1c); a higher resolution map of the same LRRK2^RCKW^ tetramer that excludes the microtubule (5.9Å) (Extended Data Fig. 3g-i); maps of pseudo-two-fold symmetry-related LRRK2^RCKW^ monomers along a filament that face either the minus (“−”) (5.2Å) or plus (“+”) (5.0Å) end of the microtubule, which revealed their different contacts with the microtubule (Fig. 1e,f and Extended Data Fig. 1c); and a consensus structure of LRRK2^RCKW^ that gave the highest resolution for the kinase domain (4.5Å) (Extended Data Fig. 1c).

The LRRK2^RCKW^ filaments are formed by two different homotypic dimer interfaces, involving either COR-B:COR-B or WD40:WD40 interactions (Fig.1c), in agreement with what had been predicted by modeling^30,31^. Each interface has a pseudo-two-fold axis of symmetry perpendicular to the microtubule axis. The ROC domain points toward and contacts the microtubule (Fig.1c-f). Interestingly, our in vitro reconstituted filaments of LRRK2^RCKW^ differ from those formed by full length LRRK2 in cells^30^, with LRRK2^RCKW^ forming a triple (rather than double) helix, with the strands packed closer together. Despite these differences, the pitch of the helix is the similar in both cases (see Methods and Supplementary Table 2).

We previously hypothesized that LRRK2’s kinase must adopt a closed conformation to form filaments around microtubules^31^. Our current structure agrees with this prediction (Fig.1g,h). To determine whether the closed conformation of the kinase was a consequence of the presence of MLi-2, which would be expected to stabilize that state, we solved a structure of microtubule-associated LRRK2^RCKW^ filaments in its absence (Extended Data Fig. 2). Although these filaments are less well ordered than those formed in the presence of MLi-2 (Extended Data Fig. 1a) and thus resulted in a lower resolution reconstruction (7.0Å), the final map still fit a closed-kinase model of LRRK2^RCKW^ better than its open form (Extended Data Fig. 3a,b). Finally, the conformation of the kinase in the microtubule-associated LRRK2^RCKW^[I2020T] filaments appears to be somewhat more closed than that predicted by AlphaFold^41,42^ for the active state of full-length LRRK2 (Extended Data Fig. 3c-f). We cannot determine at this point whether this difference is a consequence of the absence of the amino-terminal half of LRRK2, the presence of the I2020T mutation in our filaments, a small difference in the AlphaFold modeling, or a consequence of the formation of the filaments themselves.

It was previously proposed that the ROC domain would mediate binding of LRRK2 to microtubules due to its proximity to the microtubule surface in the cryo-ET map of the filaments in cells^30^. However, no density was seen connecting the ROC domain, or any other domain, to the microtubule in the cryo-ET map^30^. In contrast, our cryo-EM map showed clear density connecting LRRK2^RCKW^ and the microtubule (Fig.1e, f). As mentioned above, the polar nature of the microtubule means that the ROC domains, which would otherwise be related by a two-fold symmetry axis perpendicular to the microtubule, are in different local environments. In agreement with this, their connections to the microtubule only became apparent when LRRK2^RCKW^ monomers were refined individually (Fig. 1c and e,f, and Extended Data Fig. 1).

### LRRK2’s dimer interfaces are important for microtubule association

We next sought to examine in more detail the role played by the WD40- and COR-B-mediated dimer interfaces in LRRK2’s ability to associate with microtubules. To build a model of the LRRK2^RCKW^ filament we used rigid-body docking of individual domains from the LRRK2^RCKW^ structure (PDB: 6VNO)^31^ (Fig.2a, b). This revealed WD40:WD40 and COR-B:COR-B interfaces very similar to those seen previously with isolated WD40 domains^43^, full-length LRRK2 COR-B:COR-B dimers^35^, and LRRK2^RCKW^ dimers in the absence of microtubules^31^. However, small differences exist when domains (ROC, COR-A/B and the N-lobe of the kinase), individually fitted into our cryo-EM map of the filaments, are compared with the corresponding portion of the COR-B-mediated dimer of full-length-LRRK2^35^ (Extended Data Fig. 3g-j). It remains to be seen whether these differences are due to the absence of the N-terminal half of LRRK2 in the microtubule-associated filaments, or to small conformational changes associated with filament formation.

**Figure 2.**
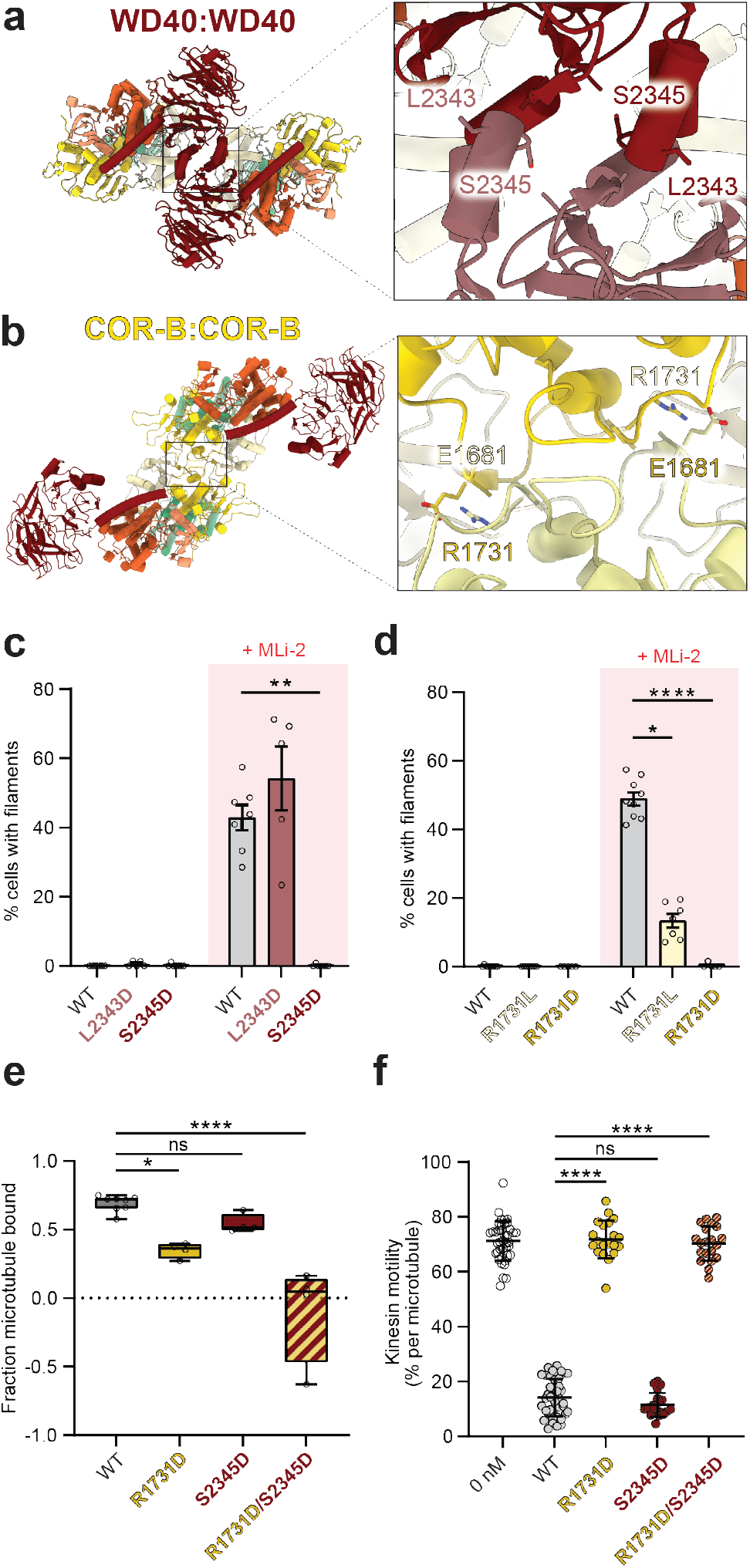
Effect of mutations in LRRK2’s WD40 and COR-B domains on filament formation and microtubule binding. **a, b**, Dimer interfaces (WD4O:WD4O and COR-B:COR-B) involved in filament formation, and location of residues tested in this work. **c**, Effect of mutations in residues in the WD40 domain (L2343D or S2345D) that have been shown to reduce dimerization of the isolated domain in vitro on the formation of MLi-2-induced filaments in cells. Cells (293T) were treated with MLi-2 (500 nM) or DMSO as a control for 2 h. Data are mean ± s.e.m. **p=0.0076, Kruskal-Wallis test with Dunn’s post hoc for multiple comparisons. **d**, Effect of mutations (R1731L/D) in a residue at the COR-B:COR-B interface on the formation of MLi-2-induced filaments in cells. Data are mean ± s.e.m. *p=0.0205, ****p<0.0001, Kruskal-Wallis test with Dunn’s post hoc for multiple comparisons. **e**, Effect of mutations in the WD40 and/or WD40 and COR-B domains on the binding of LRRK2^RCKW^ to microtubules in a pelleting assay. Tubulin concentration was 0.700 μM. Box and whisker plot center line denotes the median value, whiskers denote min and max values. *p=0.0111, ****p<0.0001, one-way ANOVA with Dunnett’s multiple comparisons test. **f**, Effect of mutations in the WD40 or WD40 and/or COR-B domains on the inhibition of kinesin motility in vitro by 50 nM LRRK2^RCKW^. Inhibition of kinesin motility was quantified as percentage of motile events per microtubule. Data are mean ± s.d. ****p<0.0001, Kruskal-Wallis test with Dunn’s post hoc for multiple comparisons.

Based on our model, we made mutants designed to disrupt both interfaces and then tested their ability to form filaments in cells and to bind microtubules or inhibit the motility of kinesin in vitro. At the WD40:WD40 interface we mutated leucine 2343 or serine 2345 to aspartic acid (L2343D or S2345D; Fig. 2a), designed to introduce a charge clash. At the COR-B:COR-B dimer interface we mutated arginine 1731 to leucine or aspartic acid (R1731L or R1731D; Fig. 2b), designed to disrupt the salt bridge with glutamic acid 1681. All mutant alleles expressed similarly to WT LRRK2 when transfected into 293T cells (Extended Data Fig. 4a,b,d,e). We also tested to each mutant for its ability to phosphorylate Rab10 in cells and found that the WD40 dimerization interface mutants had no effect on LRRK2’s kinase activity, while the COR-B dimerization interface mutants elevated it by ~2X (Extended Data Fig. 4c,f). Next, we tested the ability of these mutations to disrupt filament formation by full-length LRRK2 (WT except for the interface mutations) in cells, which is induced by MLi-2^31,31,33,29,31,31,33,34^ (Fig. 2c and Extended Data Fig. 4a,d). As previously shown, mutation of either the WD40:WD40 interface^30^ or the COR-B:COR-B interface^35^ reduced filament formation in cells. We found that S2345D, R1731L and R1731D all significantly decreased MLi-2-induced LRRK2 filament formation, with S2345D and R1731D completely abolishing our ability to detect filaments in cells (Fig. 2c, d). Surprisingly, although the L2343D mutation was previously shown to decrease dimerization in the context of a purified WD40 domain^43^, it did not reduce the formation of LRRK2 filaments in the presence of MLi-2 (Fig. 2c).

Next, we examined the effects of the mutations at the LRRK2 dimerization interfaces on LRRK2’s ability to bind microtubules or inhibit kinesin motility in vitro. To investigate LRRK2’s ability to bind microtubules in vitro, we incubated pure LRRK2^RCKW^ with in vitro assembled, taxol-stabilized microtubules and quantified the fraction of LRRK2 that pelleted with microtubules after centrifugation. While a point mutation at the WD40 dimerization interface (S2345D) did not affect LRRK2^RCKW^’s ability to pellet with microtubules, a point mutation at the COR-B interface (R1731D) reduced microtubule binding by about 50% (Fig. 2e and Extended Data Fig. 4g). Combining these mutations (R1731D/S2345D) largely abolished LRRK2^RCKW^’s interaction with microtubules (Fig. 2e and Extended Data Fig. 4g). Cryo-EM imaging of microtubules incubated with the different mutants agreed with the binding data: we observed the layer lines diagnostic of filament formation with LRRK2^RCKW^[S2345D], but not with the R1731D or R1731D/S2345D mutants (Extended Data Fig. 4h). Previously, we showed that low nanomolar concentrations of LRRK2^RCKW^ blocked the movement of dynein and kinesin motors in vitro^31^. To determine if the dimerization interfaces are required for this inhibitory effect, we monitored the motility of single GFP-tagged human kinesin-1 (“kinesin” here) molecules using total internal reflection fluorescence (TIRF) microscopy. As in the microtubule binding experiments, we found that a single point mutation at the WD40 dimerization interface (S2345D) blocked kinesin motility similarly to wild-type LRRK2^RCKW^, while a single mutant at the COR-B interface (R1731D) or the double mutant designed to disrupt both dimerization interfaces (R1731D/S2345D) no longer inhibited kinesin motility in vitro (Fig. 2f and Extended Data Fig. 4i,j). Importantly, 2D averages from cryo-EM images of LRRK2^RCKW^[R1731D/S2345D] showed that the mutations do not significantly alter the structure of the protein (Extended Data Fig. 4k).

### Electrostatic interactions drive binding of LRRK2^RCKW^ to microtubules

We next sought to test the hypothesis that binding of LRRK2 to microtubules is mediated by electrostatic interactions between the negatively charged surface of the microtubule and basic residues in LRRK2’s ROC domain. In addition to the observed charge complementarity between our model of the LRRK2 filaments and the microtubule (Fig. 3a and Deniston et al.^31^), other data support this hypothesis: (1) the symmetry mismatch between microtubules and the LRRK2 filaments suggests that there cannot be a single LRRK2-microtubule interface^30^, (2) the cryo-ET reconstruction of filaments in cells showed no clear direct contact between LRRK2 and tubulin^30^, and (3) the connections in our reconstruction only became apparent when LRRK2^RCKW^ monomers were refined individually (Fig. 1c and e,f, and Extended Data Fig. 1). To directly test this hypothesis, we developed a fluorescence-based assay to monitor binding of LRRK2^RCKW^ to microtubules in vitro. To do this, we randomly chemically labeled primary amines of LRRK2^RCKW^ with BODIPY TMR-X (“TMR”here) and used widefield fluorescence microscopy to quantify the association of TMR-LRRK2^RCKW^ with Alexa Fluor 488-labeled microtubules tethered to a coverslip. Chemical labeling did not significantly impair LRRK2^RCKW^ kinase activity as assessed by Rab8a phosphorylation in vitro (Extended Data Fig. 4l,m). In our indirect assay of filament formation, TMR-LRRK2^RCKW^ also inhibited the microtubule-based motility of kinesin (Extended Data Fig. 4n). Titration of increasing concentrations of TMR-LRRK2^RCKW^ to microtubules led to a dose-dependent increase in microtubule binding (Fig. 3b,c).Notably, LRRK2^RCKW^ binds to microtubules at low nanomolar concentrations, in agreement with our previous observations that low nanomolar concentrations of LRRK2^RCKW^ inhibit the motility of kinesin and dynein^31^. Unlabeled LRRK2^RCKW^ also binds microtubules in a bulk microtubule co-sedimentation assay (Extended Data Fig. 4o,p). To determine whether electrostatic interactions contribute to the binding of LRRK2^RCKW^ to microtubules, we tested the effect of increasing concentrations of sodium chloride on this binding. We observed a dose-dependent decrease in microtubule binding (Fig. 3d and Extended Data Fig. 4q). We also observed a salt-dependent decrease in microtubule binding for unlabeled LRRK2^RCKW^ as measured by bulk co-sedimentation with microtubules (Extended Data Fig. 4o,p).

**Figure 3.**
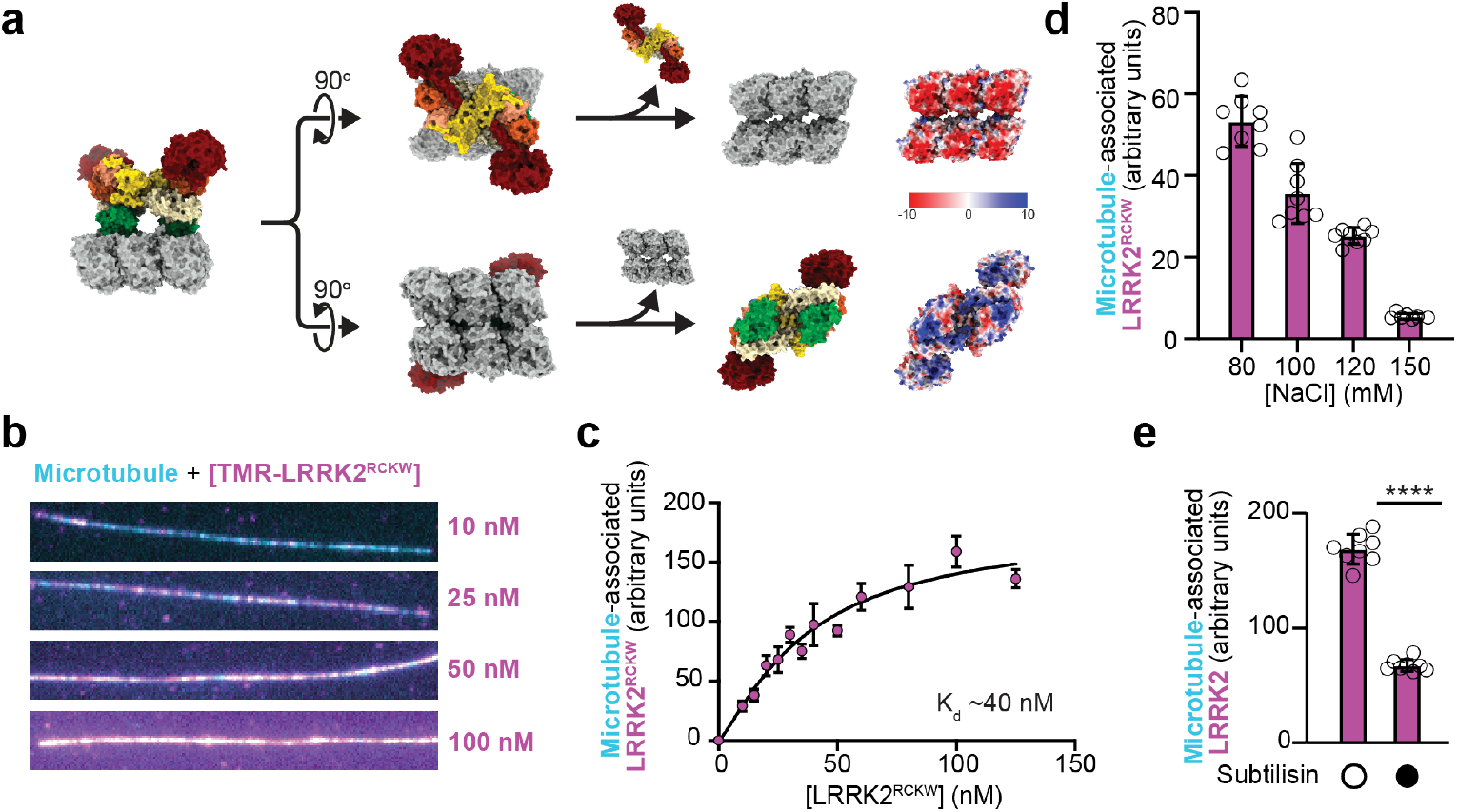
LRRK2^RCKW^ interacts with the microtubule via electrostatic interactions. **a**, Charge distribution in the molecular model for microtubule-associated LRRK2^RCKW^ filaments (Fig.1). Themodel is shown in surface representation on the left and is then split to reveal the microtubule surface facing LRRK2^RCKW^ (top) or the LRRK2^RCKW^ surface facing the microtubule (bottom). The Coulomb potential of thosesurfaces is shown on the right. The acidic C-terminal tubulin tails that further contribute negative chargedensity to the microtubule are disordered in our structure and not included here. **b**, Representative imagesof randomly labeled TMR-LRRK2^RCKW^ (magenta), bound to microtubules labeled with Alexa Fluor 488 andtethered to a coverslip (cyan). The concentrations of TMR-LRRK2^RCKW^ are indicated on the right. **c**, Quanti-fication of data represented in (b). Images were flatfield corrected, average TMR-fluorescence intensity wasmeasured along each microtubule in a given field of view, and an average value per field of view was calculated, normalized for microtubule length. Data are mean ± s.d., n=8 fields of view. **d**, Binding of 100 nMTMR-LRRK2^RCKW^ to microtubules in the presence of increasing concentrations of sodium chloride, quantifiedfrom the assay exemplified by (b). **e**, Binding of 50 nM TMR-LRRK2^RCKW^ to microtubules untreated or pre-treated with subtilisin, quantified from the assay exemplified by (b). Data are mean ± s.d., n=8 fields of view.****p<0.0001, unpaired t-test with Welch’s correction.

Tubulin carries an overall negative charge, and the disordered, negatively charged, glutamate-rich carboxy-terminal tails of tubulin are known to contribute to microtubule binding by many microtubule-associated proteins^44^. We tested the contribution of the tubulin tails to the LRRK2^RCKW^-microtubule interaction by removing them with the protease subtilisin, which cleaves tubulin near its carboxy-terminus^45^. Cleavage of tubulin tails decreased LRRK2^RCKW^’s ability to bind microtubules by ~50% (Fig. 3e and Extended Data Fig. 4r). Together, these results show that LRRK2’s interaction with the microtubule is driven by electrostatic interactions and is mediated in part by the carboxy-terminal tails of tubulin.

### LRRK1^RCKW^ adopts a similar overall fold to LRRK2^RCKW^

To hone in on specific residues in LRRK2 that might be important for mediating its interaction with microtubules, we used a comparative approach with its closest homolog, LRRK1. While LRRK2 has been linked to both familial and sporadic PD^1–6^, LRRK1 is not clinically associated with PD^46^, but instead is implicated in metabolic bone disease and osteopetrosis^47–50^. Many of LRRK1’s domains are relatively well conserved with LRRK2, with 41%, 48%, 46%, and 50% similarity between the leucine-rich repeat (LRR), ROC, COR, and kinase domains, respectively. The amino- and carboxy-termini of LRRK1 and LRRK2 are more divergent; LRRK1 lacks the amino-terminal armadillo repeats, and it is unclear based on sequence analyses whether LRRK1, like LRRK2, contains a WD40 domain, with only 27% sequence similarity in this region.

We began by solving a structure of LRRK1 using cryo-EM (Fig. 4a and Extended Data Fig. 5a). To do so, we expressed and purified the amino acids in LRRK1 that corresponded to LRRK2^RCKW^ (residues 631 to 2015; referred to as LRRK1^RCKW^; Fig. 4a). The resolution of the LRRK1^RCKW^monomer (5.8Å) was limited by the same strong preferred orientation we had observed for the LRRK2^RCKW^monomer^31^. While LRRK2^RCKW^ forms trimers, which allowed us to solve its high-resolution structure^31^, we saw no evidence of trimer formation by LRRK1^RCKW^. Our structure, obtained in the presence of GDP but in the absence of ATP, shows that LRRK1^RCKW^ adopts a similar overall J-shaped domain organization as that of LRRK2^RCKW^ and does contain a WD40 domain (Fig. 4a,b). Our map revealed that the αC helix in the N-lobe of LRRK’s kinase is ~4 turns longer than that in LRRK2 (Fig. 4c), a feature that was correctly predicted by the AlphaFold^41,42^ model of LRRK1. Our structure also revealed a density corresponding to a carboxy-terminal helix extending from the WD40 domain and lining the back of the kinase domain, as is the case for LRRK2, but the LRRK1 carboxy-terminal helix appears to be shorter (Fig. 4d). Our map disagrees with the LRRK1 structure predicted by AlphaFold, which has a longer C-terminal helix (Fig. 4d). The significance of this difference is not clear at this time as the AlphaFold structure was modeled in the active conformation (closed kinase), while our cryo-EM map of LRRK1^RCKW^ is in an inactive, open kinase conformation and lacks the N-terminal repeats. At the current resolution, LRRK2^RCKW^ and LRRK1^RCKW^ are otherwise very similar, confirming that the overall domain organization is conserved between these two proteins.

**Figure 4.**
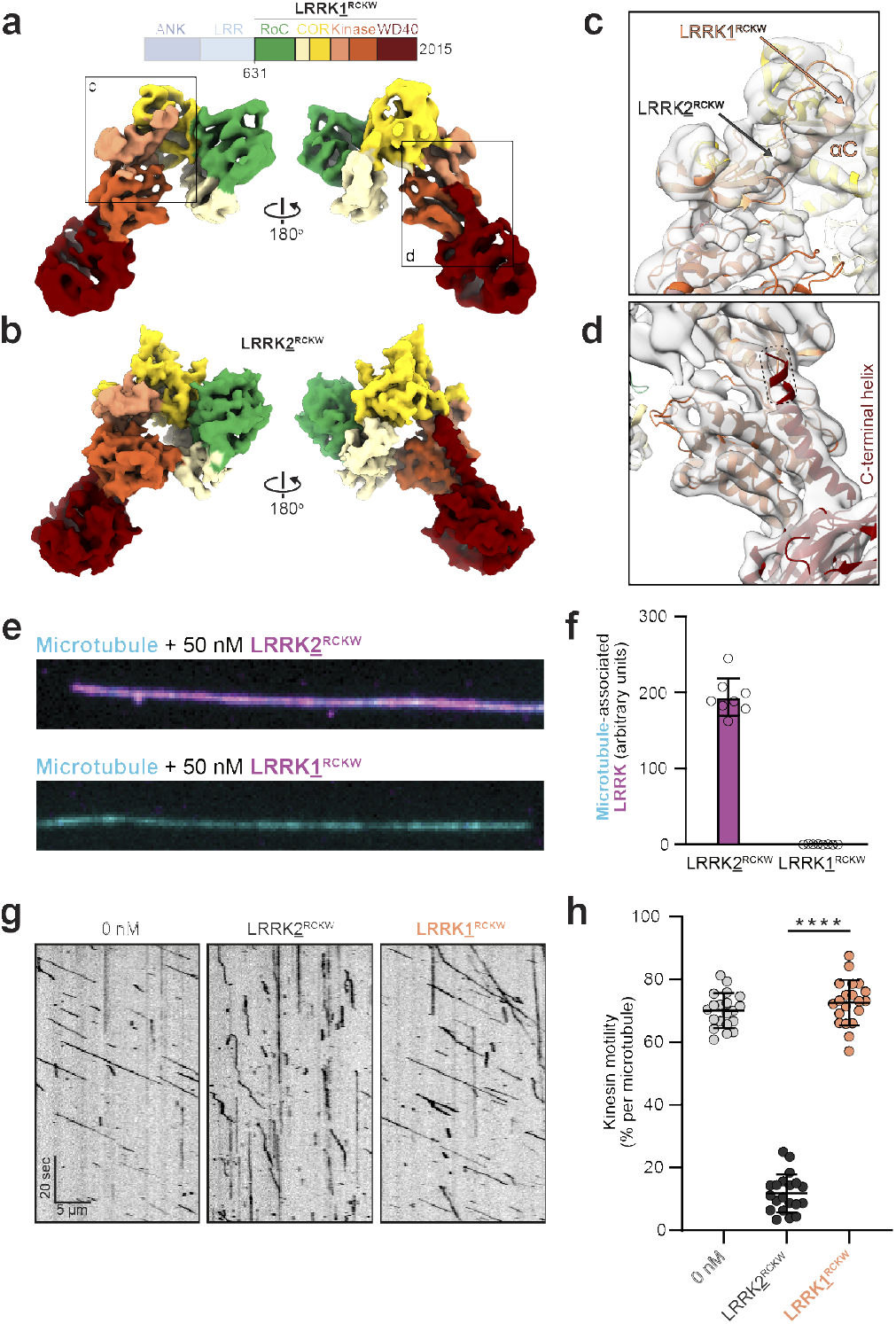
LRRK1^RCKW^ is structurally similar to LRRK2^RCKW^ but does not bind to microtubules. **a**, Cryo-EM map (5.8Å) of a LRRK1^RCKW^ monomer, with domains colored according to the scheme shown above. **b**, The molecular model for LRRK2^RCKW^ (PDB:6VNO) is shown as a calculated 6Å density (molmap command in ChimeraX), in the same orientations used for LRRK1^RCKW^ in (a). **c,d**, Close ups of the LRRK1^RCKW^ map shown in (a) with the AlphaFold model of LRRK1 docked into it. These close ups highlight the difference in length in the αC helix between LRRK1 and LRRK2 (c), and a difference between our experimental map of LRRK1^RCKW^ and the AlphaFold model of LRRK1 (d). **e**, Representative images of Alexa Fluor 488-labeled microtubules (cyan) incubated with 50nM of either LRRK2^RCKW^ (magenta, top) or LRRK1^RCKW^ (magenta, bottom). **f**, Quantification of data exemplified by (e), as outlined in Figure 3, above. Data are mean ± s.d., n=8 fields of view. **g**, Example kymographs for singlemolecule kinesin motility assays alone or in the presence of 100nM of either LRRK2^RCKW^ or LRRK1^RCKW^. **h**, Quantification of data exemplified by (g) as percentage of motile kinesin events per microtubule. Data are mean ± s.d. ****p<0.0001, Kruskal-Wallis test with Dunn’s post hoc for multiple comparisons.

### LRRK1^RCKW^ does not bind microtubules

Given the structural similarity between LRRK1 and LRRK2 (Fig. 4a,b), we wondered whether LRRK1 could also bind microtubules. To assess this, we randomly chemically labeled LRRK1^RCKW^ with BODIPY TMR-X and used widefield fluorescence microscopy to quantify microtubule-binding in vitro. We did not observe association of TMR-LRRK1^RCKW^with microtubules (Fig. 4e,f), suggesting that, unlike LRRK2^RCKW^, LRRK1^RCKW^ does not bind microtubules. As an alternative way of testing LRRK1^RCKW^’s binding to microtubules, we measured the effect of unlabeled LRRK1^RCKW^ on kinesin motility. Unlike LRRK2^RCKW^, LRRK1^RCKW^ had no effect on kinesin motility even at a concentration of 100nM, consistent with its inability to bind microtubules (Fig. 4g,h and Extended Data Fig. 5b). In addition, unlabeled LRRK1^RCKW^ did not co-sediment with microtubules (Extended Data Fig. 5c,d). Together these data show that, in contrast to LRRK2, LRRK1 does not interact with microtubules.

### Basic residues in LRRK2’s ROC domain are important for microtubule binding

Next, we sought to use our discovery that LRRK1^RCKW^ and LRRK2^RCKW^ share a similar overall structure, but only LRRK2^RCKW^ binds microtubules, to identify specific amino acids in LRRK2 that are important for microtubule binding. A sequence alignment of LRRK1 and LRRK2 revealed several basic patches in LRRK2 that are not well conserved in LRRK1 (Extended Data Fig. 5e). These basic patches create a positively charged surface on the part of LRRK2’s ROC domain facing the microtubule that is absent in LRRK1 (Fig. 5a,b). The patches correspond to residues 1356-1359 (KTKK in human LRRK2), 1383-1386 (KRKR in human LRRK2), and 1499-1502 (KLRK in human LRRK2). In our highest resolution maps, where we refined individual LRRK2^RCKW^ monomers in the filament and their contacts with the microtubule (Fig. 1e,f), the strongest density connecting LRRK2^RCKW^ to tubulin involves the 1356-1359 and 1383-1386 basic patches (Fig. 5c and Extended Data Fig. 5f). To determine if these basic patches are required for LRRK2’s interaction with microtubules, we mutated two basic residues to alanine in each patch in the context of LRRK2^RCKW^ (K1358A/K1359A or R1384A/K1385A) and tested the mutants’ ability to bind to microtubules in vitro. Both mutants showed a significant decrease in microtubule binding in a microtubule co-sedimentation assay compared with wild-type LRRK2^RCKW^ (Fig. 5d). We also tested the ability of LRRK2^RCKW^[K1358A/K1359A] to inhibit kinesin motility in vitro (Fig. 5e and Extended Data Fig. 5g,h) and found that it showed a significant reduction in its inhibition compared to wild-type LRRK2^RCKW^. Finally, we introduced full-length GFP-LRRK2 carrying either of the two basic patch mutations into human 293T cells and quantified microtubule-association in the absence or presence of MLi-2. In the absence of MLi-2, all three constructs (wild-type and the two basic patch mutants) formed little or no filaments in cells (Fig. 5f and Extended Data Fig. 5i). Treatment with MLi-2 resulted in the appearance of filaments in a significant percentage of cells carrying wildtype LRRK2 but failed to induce filament formation in cells carrying the basic patch mutants (Fig. 5f and Extended Data Fig. 5i). We also tested whether GFP-LRRK2 carrying the PD-linked I2020T mutation, which is known to result in filament formation in cells in the absence of MLi-2^29,31,31,33,34^ (Fig. 5g), is sensitive to a basic patch mutation. Indeed, GFP-LRRK2[I2020T] no longer formed microtubule-associated filaments in cells when also carrying the K1358A/K1359A mutation (Fig. 5g and Extended Data Fig. 5j). In agreement with our data on microtubule binding in vitro and filament formation in cells, cryo-EM imaging of microtubules incubated with LRRK2^RCKW^ carrying either of the two basic patch mutants did not show the layer lines diagnostic of filament formation (Extended Data Fig. 4h). Class averages from cryo-EM images of those mutants also showed that the mutations do not significantly alter the structure of the protein (Extended Data Fig. 4k).

**Figure 5.**
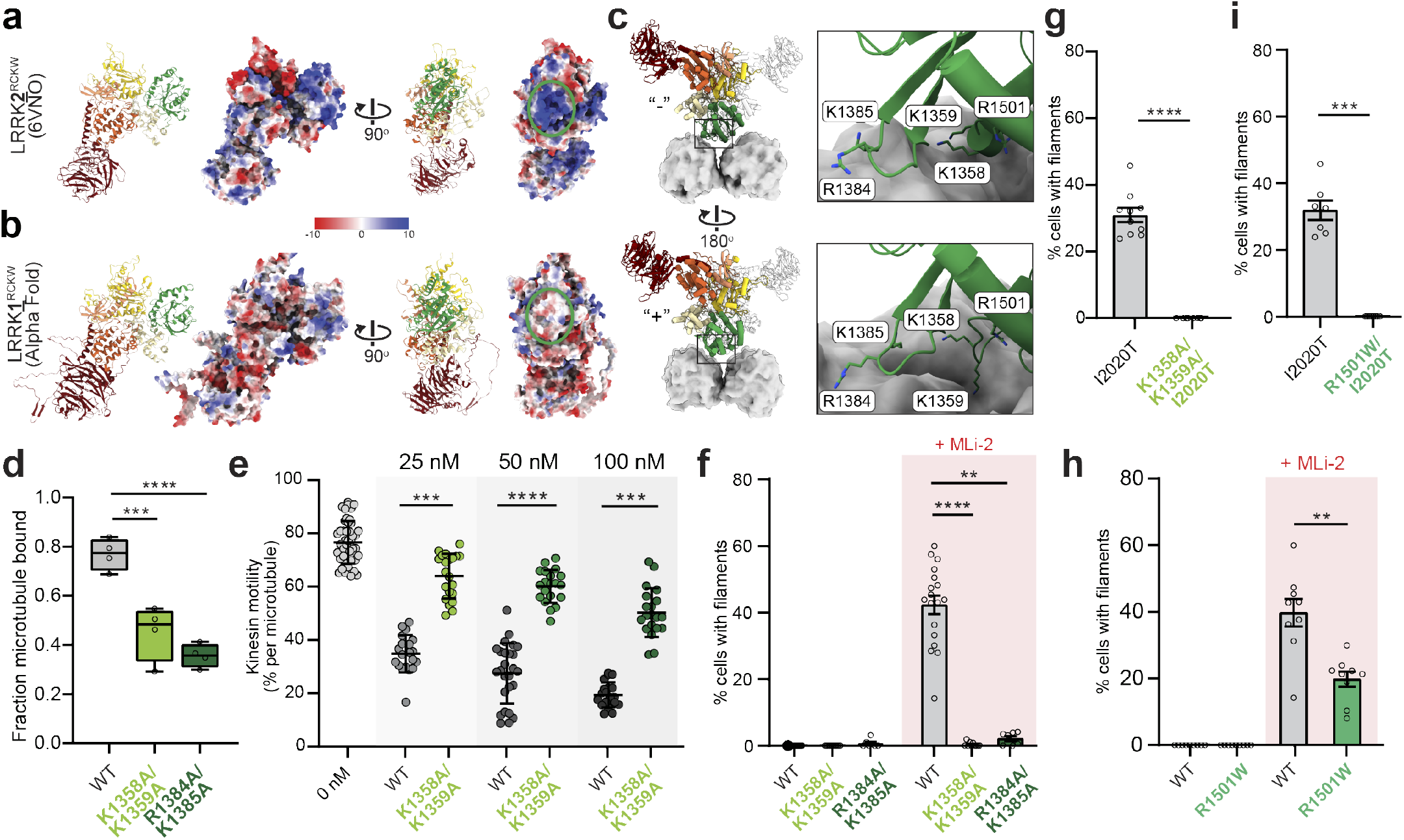
Basic patches in the ROC domain are involved in LRRK2’s binding to microtubules. **a,b**, Surface charge distribution (Coulomb potential) for LRRK2^RCKW^ (PDB:6VNO) (a) and the AlphaFold model for LRRK1 (b). The green oval on the right highlights the region in the ROC domain, which faces the microtubule in the filament structure, where basic patches are present (and conserve) in LRRK2 but absent in LRRK1. **c**, Molecular model of the microtubule-bound LRRK2^RCKW^ filament with tubulin shown in surface representation. “Top” and “bottom” show the two monomers in a dimer, along with close ups highlighting basic residues near the microtubule surface tested here. **d**, Binding of LRRK2^RCKW^, wild-type or carrying mutations in the ROC domain’s basic patches, to microtubules using a pelleting assay. Tubulin concentration was 0.432 μM. Box and whisker plot center line denotes the median value, whiskers denote min and max values.***p=0.0006, ****p<0.0001, one-way ANOVA with Dunnett’s multiple comparisons test. **e**, Single-molecule motility assays for kinesin alone or in the presence of increasing concentrations of LRRK2^RCKW^, either wild-type or carrying mutation in the ROC domain. Inhibition of kinesin motility was quantified as percentage of motile events per microtubule. Data are mean ± s.d. ***p=0.0001, ****p<0.0001, ***p=0.0002, Kruskal-Wallis test with Dunn’s post hoc for multiple comparisons. **f**, Quantification of microtubule-associated filament formation in cells for wild-type or basic patch mutant GFP-LRRK2 in the absence or presence of MLi-2. DMSO was used as a control for MLi-2. Data are mean ± s.e.m. **p=0.0022, ****p<0.0001, Kruskal-Wallis test with Dunn’s post hoc for multiple comparisons. **g**, Quantification of microtubule-associated filament formation in cells expressing GFP-LRRK2(I2020T) and basic patch mutant GFP-LRRK2[K1358A/K1359A/I2020T]. Data are mean ± s.e.m. ***p<0.0001, Mann-Whitney test. **h**, Same as for (f) for a recently identified PD-linked mutation in the ROC domain (R1501W). Data are mean ± s.e.m. **p=0.0017, Mann-Whitney test. **i**, Same as for (g) for GFP-LRRK2[I2020T] and GFP-LRRK2[R1501W/I2020T]. Data are mean ± s.e.m. ***p=0.0002, Mann-Whitney test.

While none of the most common PD-linked mutations in LRRK2 are found in these basic patch regions, the recently reported R1501W variant^51^ is found in the ROC domain facing the microtubule, near the basic patches we identified (Fig. 5c). To determine whether R1501W alters LRRK2’s interaction with microtubules we expressed GFP-LRRK2[R1501W] in 293T cells. In the presence of MLi-2, LRRK2[R1501W] showed a ~50% reduction in the fraction of cells containing microtubule-bound filaments compared to wild-type LRRK2 (Fig. 5h and Extended Data Fig. 5k). While the effect of the R1501W mutation was milder than that of the basic patch mutations in the context of wild-type LRRK2, it was as extreme as the basic patch mutants when combined with the I2020T mutation, where it also abolished filament formation (Fig. 5i and Extended Data Fig. 5j).

Importantly, none of the effects described above are due to changes in protein expression levels (Extended Data Fig. 5l) or to changes in the kinase activity of LRRK2 (Extended Data Fig. 5l,m). The basic patch mutants and R1501W all show similar levels of Rab10 phosphorylation in cells compared with wild-type LRRK2, and they do not alter the increased Rab10 phosphorylation seen in the context of I2020T (Extended Data Fig. 5l,m).

## Discussion

Here we report a structure of LRRK2^RCKW^ filaments bound to microtubules. Our structure (4.5Å in the best parts of our map) was obtained by reconstituting the filaments in vitro using pure components and is of much higher resolution than a previously reported cryo-ET structure of LRRK2 filaments bound to microtubules in cells (14Å)^30^. This new structure allowed us to build better models of the WD40:WD40 and COR-B:COR-B dimerization interfaces and it confirmed our previous proposal that filament formation by LRRK2 requires its kinase to be in a closed (active) conformation^31^. This provides a structural explanation for the observation that LRRK2-specific Type 1 kinase inhibitors, which are expected to stabilize the closed conformation of the kinase, induce filament formation in cells^30,31,33^. We also report a structure of the catalytic half of LRRK1 (LRRK1^RCKW^), LRRK2’s closest homologue. While of modest resolution (5.8Å), this structure shows that the overall fold of the ROC, COR, Kinase and WD40 domains, and their spatial arrangement, are similar in both LRRK proteins. We also undertook a comparative analysis of the two proteins with respect to microtubule interactions. We found that LRRK1 does not bind to microtubules in vitro, while LRRK2 does so with high affinity (K_d_ ~ 40 nM). As shown previously^30^, the ROC domain of LRRK2 faces the microtubule in the filament. By comparing the sequences and structures of LRRK1^RCKW^ and LRRK2^RCKW^ we identified several microtubule-facing basic patches in the ROC domain of LRRK2 that are not well conserved in LRRK1. These basic patches were in regions where we observed density connecting LRRK2^RCKW^ to the microtubule in our cryo-EM map. We discovered that mutating two basic amino acids in LRRK2’s ROC domain was sufficient to block microtubule binding both in cells and in vitro. We also showed that a PD-linked variant (R1501W)^51^, which is located in the same region of the ROC domain, decreased microtubule binding in cells. Together, this work provides important insights and tools for probing the cellular function and localization of LRRK2 and for designing LRRK2-specific kinase inhibitors.

The symmetry mismatch between the LRRK2 filaments and the microtubule, first seen in the cryo-ET reconstruction of the filaments in cells^30^, remains one of the most striking features of the structure. The previous reconstruction, at 14Å, did not show any density connecting LRRK2 to the microtubule; therefore, how the two interacted despite the symmetry mismatch remained a mystery. The higher resolution of our map, and our ability to process LRRK2^RCKW^ monomers individually, allowed us to show that LRRK2^RCKW^ monomers that are related by (pseudo)two-fold symmetry in the filament are indeed not truly symmetric and interact with the microtubule differently. This explains why those connections had not been seen before; any processing that imposes helical symmetry to the filaments would average out non-symmetric features. Understanding whether these differences in the interaction with the microtubule translate into other changes throughout LRRK2 will require higher resolutions. Higher resolutions would also allow us to determine whether filament formation alters the structure of LRRK2 relative to its soluble forms; the fact that rigid body docking of domains into our map led to small, but detectable differences with either soluble LRRK2 dimers^35^ or the AlphaFold predicted LRRK2 structure^42^ suggests this could be the case.

A major difference between our structure of microtubule bound LRRK2^RCKW^ filaments in vitro and that of LRRK2 filaments in cells was in the number and closeness of those filaments; filaments in cells were double-helical while those we presented here are triple-helical and packed closer together. Despite these differences, the helical parameters are very similar between the structures, suggesting that the underlying structure of the filaments is similar as well. The most likely explanation for the differences is the absence of the amino-terminal repeats in our structure of the LRRK2^RCKW^ filaments. Although disordered and absent from the final map, the amino-terminal half of LRRK2 was present in the filaments reconstructed in cells^30^. Placing the AlphaFold model of LRRK2 into the cryo-ET map of filaments in cells showed major clashes between the filament itself (formed by the RCKW domains) and the amino-terminal repeats (Extended Data Fig.6). Therefore, the filaments could not form unless the amino-terminal repeats were undocked from the rest of the protein, explaining why they were not visible in the cryo-ET map. Their presence, albeit in a flexible state, could explain the larger spacing, and thus lower number of helices, seen in LRRK2 vs LRRK2^RCKW^ filaments; the disordered amino-terminal repeats could act as “spacers” that prevent the filaments from packing closer together.

Although we could visualize the connections between LRRK2’s ROC domain and the microtubule in our reconstruction, its resolution is not high enough to build a model into the density. We identified key residues in this interaction by comparing, at the structural and sequence level, LRRK1^RCKW^ and LRRK2^RCKW^. The structure of LRRK1^RCKW^, although of modest resolution, revealed that the two proteins are very similar. Key differences in the charge distribution on the ROC domain facing the microtubule, and the observation that LRRK1^RCKW^ does not interact with microtubules in our assays, provided a path for identifying mutations that could abolish microtubule binding in LRRK2 without affecting its structure or kinase activity. These will be invaluable tools to probe the significance of microtubule binding by LRRK2 in relevant cell types. Finally, the general features of the filaments— their curvature and basic patches facing a negatively charged surface (the microtubule)—raise the possibility that a similar geometry could be involved in the interaction between LRRK2 and membranes.

The data we presented here suggest that LRRK2 can bind microtubules without forming filaments, which we would define as oligomers that are long enough to completely wrap around a microtubule. The data also indicate that this binding mode is likely to be the preponderant one at the low concentrations we use in our in vitro single-molecule assays. These observations stem from comparing the ability of mutants designed to disrupt dimerization interfaces (COR-B and WD40) to bind microtubules and inhibit kinesin motility in vitro, and to form filaments in cells. Any mutant that completely abolishes a dimerization interface would allow LRRK2 to form dimers (via the other interface) but would prevent the formation of longer oligomers. While mutants predicted to break either the COR-B (R1731D) or WD40 (S2345D) interfaces abolished formation of LRRK2 filaments in cells, their effects on microtubule binding in vitro were far less extreme, with R1731D resulting in a ~50% decrease and S2345D having no significant effect. It should be noted that the S2345D mutant likely does not fully disrupt the WD40 interface since it is able to form some filaments, albeit less well-ordered ones, under the high concentrations used for cryo-EM. The mutants’ affinity for microtubules correlates with their ability to inhibit kinesin motility: R1731D is unable to inhibit the motor, while S2345D inhibits motility as much as WT does. Taken together, these data suggest that small LRRK2 oligomers, as small as a dimer, could act as roadblocks for microtubule-based transport. This possibility, along with the fact that Type 1 inhibitors stabilize the conformation of LRRK2 that favors microtubule binding, should be taken into account when designing LRRK2 inhibitors.

While LRRK2 readily binds microtubules at low concentrations in vitro, whether LRRK2 binds to and/or forms filaments around microtubules in cells expressing endogenous levels of LRRK2 remains an open question. Although the only reports of LRRK2 interacting with microtubules in cells so far have been under overexpression conditions^18,29–31,34^, only a limited number of cell types have been imaged for LRRK2 localization and to our knowledge there are no reports of live-cell imaging of endogenous LRRK2. Thus, an important future goal will be to determine the localization and dynamics of LRRK2 expressed at endogenous levels in PD-relevant cell types. A recent report suggests that a noncoding *LRRK2* PD variant leads to increased *LRRK2* expression in induced-microglia^52^. In addition, *LRRK2* expression levels are elevated in a variety of immune cells in PD patients compared to age matched healthy controls^53,54^. These findings raise the possibility that increased expression of wild-type *LRRK2* could be linked to PD. Our finding that the interaction of wild-type LRRK2^RCKW^ with microtubules acts as a potent roadblock for the microtubule-based motors dynein and kinesin^31^ suggests a mechanism for how increased *LRRK2* expression levels could be detrimental for membrane trafficking. All of the membrane cargos that LRRK2 has been implicated in trafficking—including lysosomes, endo-lysosomes, autophagosomes, and mitochondria^14^—are moved by dynein and kinesin^55––57^. Elevated LRRK2 kinase activity leading to the phosphorylation of Rab GTPases is also linked to changes in membrane trafficking and specifically in the recruitment of adaptor proteins that can bind dynein and kinesin motors^58,59^. Thus, examining the effects of increased *LRRK2* expression in combination with increased LRRK2 kinase activity may be relevant for understanding the molecular basis of Parkinson’s Disease.

## Acknowledgements

We thank our funders: the Howard Hughes Medical Institute (where S.L.R-P. is an Investigator); Aligning Science Across Parkinson’s (S.L.R.-P. and A.E.L.) the Michael J. Fox Foundation (grant number 18321 to A.E.L. and S.L.R.-P.); the A.P. Giannini Foundation (postdoctoral fellowship to D.M.S.); the Molecular Biophysics Training Grant at UC San Diego (NIH Grant T32 GM008326 supported A.M.D.); the National Institutes of Health (R01GM121772 to S.L.R.-P. and R01GM107214 to A.E.L). We also thank the UC San Diego Cryo-EM Facility, the Nikon Imaging Center at UC San Diego and Eric Griffis, and the UC San Diego Physics Computing Facility for IT support.

## Author contributions

M.M. collected and processed the LRRK2 cryo-EM data. D.M.S. and M.M. collected and processed the LRRK1 cryo-EM data. Y.X.L., D.M.S., A.M.D. and M.M. purified proteins for Cryo-EM and biochemistry experiments. D.M.S. and A.M.D. performed the biochemical and single-molecule assays with the help of Y.X.L. A.M.D. performed the cellular assays. S.L.R.-P. and A.E.L. directed and supervised the research. D.M.S., M.M., A.M.D., A.E.L. and S.L.R.-P. wrote and edited the manuscript.

## Materials and Methods

### Cloning, plasmid construction, and mutagenesis

LRRK2^RCKW^ and Rab8a protein expression vectors were cloned as previously described^31^. The LRRK1 sequence was codon optimized for expression in *Spodoptera frugiperda* (Sf9) cells and synthesized by Epoch Life Science. The DNA coding for wild-type LRRK1 residues 631-2015 (LRRK1^RCKW^) was cloned through Gibson assembly into the pKL baculoviral expression vector, with an N-terminal His6-Z-tag and TEV protease cleavage site. LRRK2 mutants were cloned using QuikChange site-directed mutagenesis (Agilent), or Q5 site-directed mutagenesis (New England Biolabs) following manufacturer’s instructions. As previously described for LRRK2^RCKW,31^, the LRRK1^RCKW^ plasmid was used for the generation of recombinant baculoviruses according to bac-to-bac expression system protocols (Invitrogen).

For mammalian expression, GFP-LRRK2 was cloned into the pDEST53 vector (Addgene 25044) as previously described^31^. LRRK2 mutants were cloned using QuikChange site-directed mutagenesis (Agilent) using standard protocols, with the exception that liquid cultures of *E. coli* were grown at 30°C. EGFP-Rab10^60^ was obtained from Addgene (#49472) and pET17b-Kif5b(1-560)-GFP-His^61^ was obtained from Addgene (#15219).

### LRRK2^RCKW^ and LRRK1^RCKW^ expression and purification

N-terminally His_6_-Z-tagged LRRK2^RCKW^ was expressed in Sf9 insect cells and purified as previously described^31^. Briefly, ~1L insect cells were infected with baculovirus and grown at 27°C for 3 days. Pelleted Sf9 cells were resuspended in lysis buffer (50 mM HEPES pH 7.4, 500 mM NaCl, 20 mM imidazole, 0.5 mM TCEP, 5% glycerol, 5 mM MgCl_2_, 20 μM GDP, 0.5 mM Pefabloc, and protease inhibitor cocktail tablets) and lysed by Dounce homogenization. Clarified lysate was incubated with Ni-NTA agarose beads (Qiagen), extensively washed with lysis buffer, and eluted in buffer containing 300 mM imidazole. Protein eluate was diluted 2-fold in buffer containing no NaCl, loaded onto an SP Sepharose column, and eluted with a 250 mM to 2.5 M NaCl gradient. Protein was cleaved by TEV protease overnight. Cleaved protein was isolated by running over a second Ni-NTA column. Protein was concentrated and run on an S200 gel filtration column equilibrated in storage buffer (20 mM HEPES pH 7.4, 700 mM NaCl, 0.5 mM TCEP, 5% glycerol, 2.5 mM MgCl_2_, 20 μM GDP). Final protein was concentrated to ~20-30 μM as estimated by absorbance at 280 nm using an extinction coefficient of 140150 M^−1^cm^−1^.

### Purification of molecular motors

Human KIF5B^1-560^(K560)-GFP was purified from *E. coli* using an adapted protocol previously described (Nicholas et al. 2014). All protein purification steps were performed at 4°C unless otherwise noted. pET17b-Kif5b(1-560)-GFP-His was transformed into BL-21[DE3]RIPL cells (New England Biolabs) until an optical density at 600 nm of 0.6-0.8 and expression was induced with 0.5 mM isopropyl-β-d-thiogalactoside (IPTG) for 16h at 18°C. Frozen pellets from 7.5 liters of culture were resuspended in 120 ml lysis buffer (50 mM Tris, 300 mM NaCl, 5 mM MgCl_2_, 0.2 M sucrose, 1 mM dithiothreitol (DTT), 0.1 mM Mg-ATP, and 0.5 mM Pefabloc, pH 7.5) supplemented with one cOmplete EDTA-free protease inhibitor cocktail tablet (Roche) per 50 ml and 1 mg/ml lysozyme. The resuspension was incubated on ice for 30 min and lysed by sonication. Sonicate was supplied with 0.5 mM PMSF and clarified by centrifugation at 40,000 rcf for 60 min in a Type 70 Ti rotor (Beckman). The clarified supernatant was incubated with 15 ml Ni-NTA agarose (Qiagen) and rotated in a nutator for 1 h. The mixture was washed with 100 ml wash buffer (50 mM Tris, 300 mM NaCl, 5 mM MgCl_2_, 0.2 M sucrose, 10 mM imidazole, 1 mM dithiothreitol (DTT), 0.1 mM Mg-ATP, and 0.5 mM Pefabloc, pH 7.5) by gravity flow. Beads were resuspended in elution buffer (50 mM Tris, 300 mM NaCl, 5 mM MgCl_2_, 0.2 M sucrose, 250 mM imidazole, 0.1 mM Mg-ATP, and 5 mM βME, pH 8.0), incubated for 5 min, and eluted stepwise in 0.5 mL increments. Peak fractions were combined, and buffer exchanged on a PD-10 desalting column (GE Healthcare) equilibrated in storage buffer (80 mM PIPES, 2 mM MgCl_2_, 1 mM EGTA, 0.2 M sucrose, 1 mM DTT, and 0.1 mM Mg-ATP, pH 7.0). Peak fractions of motor solution were either flash-frozen at −80°C until further use or immediately subjected to microtubule bind and release purification. A total of 1 ml motor solution was incubated with 1 mM AMP-PNP and 20 μM taxol on ice in the dark for 5 min and subsequently warmed to room temperature. For microtubule bind and release, polymerized bovine brain tubulin was centrifuged through a glycerol cushion (80 mM PIPES, 2 mM MgCl_2_, 1 mM EGTA, and 60% glycerol (v/v) with 20 μM taxol and 1 mM DTT) and resuspended as previously described^62^ incubated with the motor solution in the dark for 15 min at room temperature. The motor-microtubule mixture was laid on top of a glycerol cushion and centrifuged in a TLA120.2 rotor at 80,000 rpm (278088*g)* for 12 min at room temperature. Final pellet (kinesin-bound microtubules) was washed with BRB80 (80 mM PIPES, 2 mM MgCl_2_, and 1 mM EGTA, pH 7.0) and incubated in 100 μl of release buffer (80 mM PIPES, 2 mM MgCl_2_, 1 mM EGTA, and 300 mM KCl, pH 7 with 7.5 mM Mg-ATP) for 5 min at room temperature. The kinesin release solution was spun at 72,000 rpm (225252*g*) in TLA100 for 7 minutes at room temperature. The supernatant containing released kinesin was supplied with 660 mM sucrose and flash frozen. A typical kinesin prep in the lab yielded 0.5 to 1.5 μM K560-GFP dimer.

### Rab8a expression and purification

Rab8a was expressed and purified as previously described^31^. Briefly, N-terminally His6-ZZ tagged Rab8a was expressed in BL21(DE3) *E. coli* cells by addition of 0.5 mM IPTG for 18 hours at 18°C. Cells were pelleted, resuspended in lysis buffer (50 mM HEPES pH 7.4, 200 mM NaCl, 2 mM DTT, 10% glycerol, 5 mM MgCl_2_, 0.5 mM Pefabloc, and protease inhibitor cocktail tablets), and lysed by sonication on ice. Clarified lysate was incubated with Ni-NTA agarose (Qiagen). Protein was washed with wash buffer (50 mM HEPES pH 7.4, 150 mM NaCl, 2 mM DTT, 10% glycerol, 5 mM MgCl_2_) and eluted in buffer containing 300 mM imidazole. Eluate was incubated with IgG Sepharose 6 fast flow beads. Following further washing, Rab8a was cleaved off IgG Sepharose beads by incubation with TEV protease at 4°C overnight. Cleaved Rab8a was isolated by incubation with Ni-NTA agarose beads followed by washing with buffer containing 25 mM imidazole. Purified Rab8a was run on an S200 gel filtration column equilibrated in S200 buffer (50 mM HEPES pH 7.4, 200 mM NaCl, 2 mM DTT, 1% glycerol, 5 mM MgCl_2_). Protein was then concentrated and exchanged into 10% glycerol for storage.

### Cryo-electron microscopy: sample preparation and imaging of filaments

LRRK2^RCKW^ filaments were prepared as previously described^31^ with the exception that 10% glycerol was used instead of 10% DMSO in all the samples save for the one that led to the initial data set (“19dec14f’), as glycerol promotes the formation of 11 and 12-protofilament microtubules. For “+MLi-2” samples, we added MLi-2 to LRRK2^RCKW^ to a final concentration of 5μM after incubating them with tubulin. The updated protocol is also available at protocols.io (http://dx.doi.org/10.17504/protocols.io.bpnrmmd6).

Cryo-EM data were collected on a Talos Arctica (FEI) operated at 200 kV, equipped with a K2 Summit direct electron detector (Gatan). Automated data collection was performed using Leginon^63^ with a custom-made plug-in to automate the targeting to areas of the sample that contained LRRK2^RCKW^ filaments. The only exception was the first data set (“19dec14f’), which was collected using Leginon’s regular raster target finder. The “19dec14f” dataset was subsequently used for training the machine learning component of the custom-made plug-in used for all other datasets. The code for the plug-in is available at (https://github.com/matyszm/filfinder).

The “Apo” reconstruction was obtained using two datasets: 836 micrographs from “19dec14f’ and 1010 micrographs from “20aug12b”. The “MLi-2” reconstruction was also obtained from two datasets: 926 micrographs from “20sep10b” and 1430 micrographs from “20sep30c”. Final micrograph counts only include micrographs with at least one usable LRRK2^RCKW^ filament. The dose per dataset varied between 5 and 5.5 electrons Å–2 s–1. To accommodate for that range, we varied the exposure time between 10 and 11 seconds, with 200-ms frames, for a total number of frames between 50 and 55, and a total dose of 55 electrons Å–2. The images were collected at the nominal magnification of 36,000x, resulting in an object pixel size of 1.16 Å. The defocus was set to −1.5 μm, with a final range of defoci from −0.5 to −2.5 μm due to the nature of the lacey carbon grids and the collection strategy used. All datasets are available on EMPIAR (Supplementary Table 1).

### Cryo-electron microscopy: reconstruction of LRRK2^RCKW^ bound to a microtubule

Movie frames were aligned using UCSF MotionCor2^64^ with the dose-weighting option on. CTF estimation was done with CTFFIND4^65^ using the non-dose-weighted aligned micrographs. All micrographs containing filaments were kept regardless of the CTF estimated resolution. Data processing up to the symmetry expansion step is detailed in the protocol available in protocols.io (http://dx.doi.org/10.17504/protocols.io.bwwnpfde). In brief, manual selected filaments from a subset of micrographs were 2D classified (Relion 3.1)^66^, with the best classes then acting as a reference for automated filament picking (Relion 3.1). The separation distance of the particles was set to 30 Å, which ensures each particle contains one new LRRK2^RCKW^ dimer per strand. These particles were then filtered first by classifying based on whether a microtubule is present, then followed up by another 2D classification focusing on the presence of ordered LRRK2^RCKW^ filaments if MLi-2 was present, or a blurred, disordered layer if working with apo filaments. The selected particles were then 3D classified into 6 classes (Relion 3.1), each one corresponding to a specifically sized microtubule (from 11 to 16-protofilaments). This step is inspired by MiRP^67^ and used their provided reference scaled to the appropriate pixel size. Filaments with MLi-2 present tend to favor 11-protofilament microtubules, while the apo filaments favored larger sizes. We kept all the 11-protofilament microtubules for the MLi-2 dataset and all the 12-protofilament microtubules for the apo dataset.

In order to more accurately reconstruct the LRRK2^RCKW^ filaments, the microtubule had to be digitally subtracted from the particles. To accomplish this, we refined the structure of the microtubule for each dataset (Relion 3.1) and subtracted it from the particles (Relion 3.1, using legacy subtraction mode). This allowed us to 2D classify (Relion 3.1) focusing on LRRK2^RCKW^ filaments. Particles falling into ordered 2D classes were further 3D classified (Relion 3.1). The initial reference for each subgroup (with or without MLi-2) was always a featureless cylinder and was initialized with the helical symmetry reported for microtubule-associated LRRK2 filaments in cells^30^. Subsequent rounds used the output as the reference and were allowed to refine the symmetry, often showing multiple classes (Extended Data Fig. 1b and Supplementary Table 2). Once the symmetry was found, a local refinement was done with the original un-subtracted particles to give us a LRRK2^RCKW^ filament containing some of the original microtubule density. Since our LRRK2^RCKW^ filaments each have three strands, we used symmetry expansion to extract an individual dimer from each strand. We centered the new particles on the subtraction mask and decreased the box size to 300 pixels while keeping the Å/px scale the same. This step was performed with the new subtraction function in Relion 3.1. This resulted in 206,649 particles for the MLi-2 dataset and 49,629 particles for the apo dataset. See Extended Data Figs. 1b,c and 2 for the data processing workflow. A more detailed protocol can be found in protocols.io (http://dx.doi.org/10.17504/protocols.io.bwwnpfde). After symmetry expansion, the newly generated particles were exclusively processed in CryoSPARC version 3.2.0. The first step was always to align the particles to the centered subtraction mask in order to align the particles to each other. For the particles coming from the MLi-2 dataset we were able to compare the Psi Euler angle to the original angle assigned during the microtubule only alignment. Since the particles were allowed to be flipped during the LRRK2^RCKW^ refinement, only particles showing 0+/−20 degrees and 180+/− 20 degrees were kept. Particles with a ~180 flip were flipped back to align them to the microtubule. This left us with 133,246 particles for the MLi-2 dataset. This step was skipped for the apo particles due to the lower particle count.

The MLi-2 dataset was processed in two different ways, resulting in different levels of detail in either the kinase or ROC regions. The first processing strategy was designed to achieve a better kinase reconstruction. Here, we allowed the filtered particles to be freely aligned again, ignoring the microtubule orientation. This was followed by two local refinements: the first focused on a LRRK2^RCKW^ tetramer, and the second on a single LRRK2^RCKW^ monomer. The second strategy was designed to better resolve the contacts between the ROC domain and the microtubule. Here, we only performed local refinements on the particles with the fixed microtubule orientation. To make sure the microtubule was properly aligned, we performed a local refinement focusing only on the microtubule, which resulted in a map with no ambiguity in the tubulin orientation. We then did a local refinement focused on a LRRK2^RCKW^ dimer, followed by a 3D Variability Analysis (3DVA, in cryoSPARC)^68^ focused on a single LRRK2^RCKW^ monomer with the goal of being able to separate particles that have intact LRRK2^RCKW^ in them. We analyzed the components generated and determined that components 1 and 2 ranged from a well formed LRRK2^RCKW^ to having discontinuities or weak densities. We only kept particles with a negative value for at least one of the two components. Following that, we did another refinement for a LRRK2^RCKW^ dimer using the filtered particles, and then used a smaller mask to cover either the “+” or “−” LRRK2^RCKW^ (see Extended Data Fig. 1c) along with a small part of the microtubule. This resulted in maps showing the ROC domains interacting with the microtubule.

For the apo sample, only the freely aligned approach was used as the particle count and resolution was too low to filter the particles by microtubule orientation. After freely aligning the particles to the recentered subtraction mask, we performed a local refinement focused on the LRRK2^RCKW^ tetramer. To help the alignment, we used a 20 Å low-passed LRRK2^RCKW^ tetramer reference built by rigid body fitting 4 copies of LRRK2^RCKW^ into the early 9 Å reconstruction. This new reconstruction was still noisy, most likely due to multiple conformations being present. While Relion Class3D did not work on this dataset, we were able to use 3DVA again to help us find a component to separate apo LRRK2^RCKW^ into classes. Component 1 resulted in more a detailed reconstruction at both positive and negative ends of the spectra than the starting structure. We reconstructed both sets, and while both were able to reach ~7 Å resolution, the data with the positive component 1 resulted in a more continuous map and was chosen as the final map.

### Model creation and refinement: LRRK2 ROC domain interacting with the microtubule

For modeling we used the maps where the refinement had been focused on the interactions of the ROC domain with the microtubule, facing either towards the plus or minus end of the microtubule. For the initial model, we used LRRK2’s AlphaFold^41,42^ model (Q5S007) as it had the most complete loops available for the ROC domain (using residues 1332-1525). Since AlphaFold models lack ligands, we added GDP based on the placement in previous structures^31,35^. Because the ROC domain only occupies a small portion of the map and some microtubule density is present, we added tubulin dimers (PDB code: 1TUB) to provide a restraint during refinement. Initial refinement was done using Rosetta (ver 3.13) and Frank DiMaio’s cryo-EM refinement scripts. 200 models were generated from each map. Tubulin dimers were removed from the model before further quantification. Models with the best energy score and fit to the density were manually inspected. Small modeling errors were corrected in Isolde by hand and refined one more time in Rosetta using Relax with the map density loaded in as a restraint. 5 models were selected for each map and converted to polyalanine models except for residues of interest (K1358, K1359, R1384, K1385, R1501).

### Cryo-electron microscopy: sample preparation and imaging of LRRK1^RCKW^

The protocol for preparing LRRK1^RCKW^ grids is available at protocols.io (http://dx.doi.org/10.17504/protocols.io.b3rqqm5w). Briefly, the protein was spun down after thawing, and kept on ice until grid making. We used UltrAuFoil Holey Gold 1.2/1.3 300 mesh grids and plasma cleaned them in a Solarus II (Gatan) using the QuantiFoil Au preset. Immediately before freezing, LRRK1^RCKW^ was added to “LRRK2 buffer” (20 mM HEPES pH 7.4, 80 mM NaCl, 0.5 mM TCEP, 2.5 mM MgCl_2_, 20 μM GDP) to the desired concentration (2-6 μM protein). We used a Vitrobot Mark IV (FEI) to freeze our samples.

Cryo-EM data were collected on a Talos Arctica (FEI) operated at 200 kV, equipped with a K2 Summit direct electron detector (Gatan). Automated data collection was performed using Leginon.^63^ Reconstruction was done with 4 datasets (“19dec11a”: 847 micrographs, “19dec21c”:926 micrographs, “20sep11a”: 904 micrographs, and “21jan18d”: 952 micrographs). One of the datasets (“20sep11a”) was collected at a 20° tilt. The exposure of the micrographs varied to achieve a total dose of 55 electrons Å–2. The images were collected at a nominal magnification of 36,000x, resulting in an object pixel size of 1.16 Å. The defocus was set to −1.5 μm, which gave a range of defoci of −0.8 to −1.8 μm over all datasets. All datasets are available on EMPIAR (Supplementary Table 1).

### Cryo-electron microscopy: reconstruction of LRRK1^RCKW^

Movie frames were aligned in cryoSPARC using the “patch motion correction’’ program. CTF estimation was also done in cryoSPARC using the “patch CTF estimation” program. Images were manually screened for any obvious defects and removed from further processing if defects were found. Particle picking was done with a mixture of a crYOLO^69^ set previously trained for LRRK2^RCKW,31^ and simple blob picking followed by a round of 2D classification to remove obvious contaminants. Both methods gave similar results, and both were used depending on whether the picking was done on the fly (blob picker) or later (crYOLO). The final particle count was 645,743.

At this point, 2D classification was used on the combined particles. Only classes showing an intact RCKW-like shape were kept. Using ab-initio reconstruction gave us 2 classes, with 2/3 of the particles ending in the intact class. We recovered additional intact particles from the broken class after another round of 2D classification. Combining class 1 and the good 2D classes gave us 131,821 particles, from which we were able to obtain a 5.8 Å map with some stretched features, likely due to preferred orientation. To lower the impact of preferred orientation, we used “Rebalance 2D” with the rebalance factor set to 0.7, making sure the smallest supergroup is at least 70% of the size of the largest. While the resolution dropped to 6.5 Å, the severity of the stretching was reduced.

Despite this improvement, the map contained discontinuous density on the edges of the mask, suggesting problems with the automatically generated mask. We remade the mask by basing it on homology models of LRRK1^RCKW^ domains (ROC, COR, and Kinase; SWISS model^70^,) and LRRK2’s WD40 domain that we rigid body fitted into the current best LRRK1^RCKW^ density and used molmap in ChimeraX^71^ to create a map to serve as the mask. This map was low-passed to 15 Å, dilated by 8 px, and soft padded by another 8 px. This was then used to refine the structure one more time. This new map still contained artifacts in the ROC and COR-A region. We used 3DVA to analyze the structure and found a component showing slight movement of these domains. We selected to focus on particles in the more “closed” state. Refining these new particles gave us a better-defined map without artifacts at 5.8 Å resolution after using cryoSPARC’s Non-Uniform Refinement.

### Single-molecule microscopy and motility assays

Single-molecule kinesin motility assays were performed as previously described^31^. Imaging was performed with an inverted microscope (Ti-E Eclipse; Nikon) equipped with a 100x 1.49 NA oil immersion objective (Plano Apo; Nikon). The microscope was equipped with a LU-NV laser launch (Nikon), with 405 nm, 488 nm, 532 nm, 561 nm, and 640 nm laser lines. The excitation and emission paths were filtered using appropriate single bandpass filter cubes (Chroma). The emitted signals were detected using an electron multiplying CCD camera (Andor Technology, iXon Ultra 888). The xy position of the stage was controlled by ProScan linear motor stage controller (Prior). Illumination and image acquisition were controlled by NIS Elements Advanced Research software (Nikon).

Single-molecule motility assays were performed in flow chambers assembled as previously described^72^. Biotin-PEG-functionalized coverslips (Microsurfaces) were adhered to glass slides using double-sided scotch tape. Each slide contained four flow-chambers. Taxol-stabilized microtubules (approximately 15 mg ml^−1^) with 10% biotin-tubulin and 10% Alexa 405-tubulin were prepared as previously described^72^. For each motility experiment, 1 mg ml^−1^ streptavidin (in 30 mM HEPES, 2 mM magnesium acetate, 1 mM EGTA, 10% glycerol) was incubated in the flow chamber for 3 min. A 1:150 dilution of taxol-stabilized microtubules in motility assay buffer (30 mM HEPES, 50 mM potassium acetate, 2 mM magnesium acetate, 1 mM EGTA, 10% glycerol, 1 mM DTT and 20 μM Taxol, pH 7.4) was added to the flow chamber for 3 min to adhere polymerized microtubules to the coverslip. Flow chambers containing adhered microtubules were washed twice with LRRK2 buffer (20 mM HEPES pH 7.4, 80 mM NaCl, 0.5 mM TCEP, 5% glycerol, 2.5 mM MgCl_2_ and 20 μM GDP). Flow chambers were then incubated for 5 min with either LRRK2 buffer alone or LRRK2 buffer containing the indicated concentration of wildtype or mutant LRRK2^RCKW^. Before the addition of kinesin motors, the flow chambers were washed three times with motility assay buffer containing 1 mg ml^−1^ casein. The final imaging buffer for motors contained motility assay buffer supplemented with 71.5 mM βME, 1 mM Mg-ATP, and an oxygen scavenger system, 0.4% glucose, 45 μg/ml glucose catalase (Sigma-Aldrich), and 1.15 mg/ml glucose oxidase (Sigma-Aldrich). The final concentration of kinesin in the motility chamber was 1 nM. K560-GFP was imaged every 500 ms for 2 min with 25% laser (488) power at 150 ms exposure time. Each sample was imaged no longer than 15 min. Each technical replicate consisted of movies from at least two fields of view containing between 5 and 10 microtubules each.

### Single-molecule motility assay analysis

Kymographs were generated from motility movies using ImageJ macros as described previously^72^. Specifically, maximum-intensity projections were generated from time-lapse sequences to define the trajectory of particles on a single microtubule. The segmented line tool was used to trace the trajectories and map them onto the original video sequence, which was subsequently re-sliced to generate a kymograph. Brightness and contrast were adjusted in ImageJ for all videos and kymographs. Motile and immotile events (>1 s) were manually traced using ImageJ and quantified for run lengths and percent motility. Run-length measurements were calculated from motile events only. For percent motility per microtubule measurements, motile events (>1 s and >785 nm) were divided by total events per kymograph. Bright aggregates, which were less than 5% of the population, were excluded from the analysis. Data visualization and statistical analyses were performed in GraphPad Prism (9.2; GraphPad Software) and ImageJ (2.0).

### Microtubule sedimentation binding assay

Porcine brain tubulin was purchased from Cytoskeleton, Inc. Taxol-stabilized microtubules were polymerized at a final concentration of ~ 2.5 mg/mL, and free tubulin was removed by ultracentrifugation at 108628 x g for 15 min at 37°C through a 64% glycerol cushion. The resulting microtubule pellet was resuspended in LRRK2 binding buffer (20 mM HEPES pH 7.4, 110 mM NaCl, 0.5 mM TCEP, 5% glycerol, 2.5 mM MgCl_2_, 20 μM GDP and 20 μM Taxol). Tubulin concentration was determined by comparison of the polymerized microtubule stock to actin standards on SDS-PAGE.

For a typical LRRK^RCKW^ microtubule cosedimentation assay, 200 nM LRRK^RCKW^ was incubated at room temperature for 10 minutes with varied concentrations of microtubules in buffer containing 20 mM Hepes pH 7.4, 110 mM NaCl, 0.5 mM MgCl_2_, 0.5 mM TCEP, 5% glycerol, 20 μM GDP, 20 μM taxol. Microtubules were then pelleted by ultracentrifugation (15 minutes, 108628 g, 25 degrees). To quantify the depletion of LRRK2^RCKW^, samples of the supernatant were taken and boiled for 10 min in SDS buffer. Samples were run on 4-12% polyacrylamide gels (NuPage, Invitrogen) and stained with SYPRO-Red Protein Gel Stain (ThermoFisher) for protein detection. Binding curves were fit in GraphPad Prism (9.2; GraphPad Software) with a nonlinear regression hyperbolic curve.

### TMR labeling

BODIPY TMR-X NHS Ester (ThermoFisher) was used to fluorescently label LRRK2^RCKW^ and LRRK1^RCKW^. For a typical 40 uL labeling reaction, dye was added at a ratio of 1:1 to ~20 uM LRRK2^RCKW^, followed by incubation at room temperature for 1 hour. Excess dye was removed by two consecutive buffer exchanges through Micro Bio-Spin P-6 desalting columns (Bio-Rad). Protein concentration and labeling efficiency were estimated using a NanoDrop Microvolume Spectrophotometer.

### Widefield fluorescence microtubule binding assay

Imaging was performed with an inverted microscope (Nikon, Ti-E Eclipse) equipped as described above (single-molecule microscopy and motility assays).

LRRK2^RCKW^ microtubule-binding experiments were performed in flow chambers made as described above (single-molecule microscopy and motility assays). LRRK^RCKW^ was labeled with TMR (TMR labeling, above); taxol-stabilized microtubules were polymerized from a mixture of unmodified, biotinylated, and Alexa-488 labeled bovine tubulin, as previously described (REF). To attach microtubules to the coverslip, flow chambers were incubated with 0.5 mg/mL streptavidin for 3 minutes, washed twice in buffer (30 mM Hepes pH7.4, 50 mM KOAc, 2 mM MgOAc, 1 mM EGTA, 10% glyercol, 1 mM DTT, and 0.2 mM taxol), and then incubated with microtubules for 3 minutes. Microtubules were washed twice in buffer (20 mM Hepes pH 7.4, 80 mM NaCl, 0.5 mM MgCl_2_, 0.5 mM TCEP, 5% glycerol, 20 μM GDP), and then incubated with varied concentrations of LRRK2^RCKW^ (6.25 nM – 50 nM) for 5 minutes. Multiple fields of view were imaged along the flow chamber with the objective in widefield illumination, with successive excitation at 488 nm (15% laser power, 100 ms exposure) and 561 nm (25% laser power, 100 ms exposure).

Image analysis was performed with ImageJ. Average TMR-LRRK2^RCKW^ fluorescence intensity per microtubule was calculated from a 1 pixel-wide line drawn along the long axis of the microtubule; overall average background fluorescence intensity was subtracted. These background-subtracted intensities were averaged over all microtubules per field of view, normalized by microtubule length, to yield a single data point. Eight fields of view at each concentration of LRRK2^RKCW^ were then averaged.

### In vitro Rab8a phosphorylation

LRRK kinase assays were performed as previously described^31^ with LRRK^RCKW^ and Rab8a purified as described above. For a typical kinase reaction, 38 nM LRRK^RCKW^ was incubated with 3.8 μM Rab8a for 30 minutes at 30 degrees in buffer containing 50 mM Hepes pH 7.4, 80 mM NaCl, 10 mM MgCl2, 1 mM ATP, 200 uM GDP, 0.5 mM TCEP. Phosphorylation of Rab8a at residue T72 by LRRK^RCKW^ was monitored by western blot using a commercially available antibody (Abcam antibody MJF-R20) as previously described^31,73^.

### Immunofluorescence, confocal microscopy, and image analysis

LRRK2 filament assays were performed as previously described^31^. Briefly, cells were plated on fibronectin-coated glass coverslips and grown for 24 h before transfection with PEI. Cells were transfected with 500 ng of indicated GFP-LRRK2 plasmids. After 24-48 h, cells were incubated at 37°C with DMSO or MLi-2 (500 nM) for 2 h. Stocks of the kinase inhibitor MLi-2 (10 mM; Tocris) were stored in DMSO at −20°C.

Cells were rinsed briefly with ice-cold 1× PBS on ice, the fixed with ice-cold 4% PFA, 90% methanol, 5 MM sodium bicarbonate for 10 min at − 20°C. Coverslips were subsequently washed three times with ice-cold PBS and then incubated with blocking buffer (1% BSA, 5% normal goat serum, 0.3% Triton X-100 in 1× PBS) for 1 h at room temperature. Primary antibodies were diluted in antibody dilution buffer (1% BSA, 0.1% Triton X100 in 1× PBS) and incubated at 4°C overnight. The following day, coverslips were washed three times with 1x PBS and incubated with secondary antibodies diluted in antibody dilution buffer for 1 h at room temperature. After secondary incubation, coverslips were washed three times with 1x PBS. Cells were briefly rinsed in ddH_2_O and mounted on glass slides using CitiFluor AF-1 mounting media (TedPella). Coverslips were sealed with nail polish and stored at 4°C. Antibodies used for immunofluorescence were used at a 1:500 dilution and included: chicken anti-GFP (Aves Labs) and goat anti-chicken-Alexa 488 (ThermoFisher). DAPI was used at 1:5000 according to the manufacturer’s recommendation (ThermoFisher).

For the LRRK2 filament analysis, experimenters were blinded to conditions for both the imaging acquisition and analysis. Cells were imaged using a Yokogawa W1 confocal scanhead mounted to a Nikon Ti2 microscope with an Apo 60x 1.49 NA objective. The microscope was run with NIS Elements using the 488nm and 405nm lines of a six-line (405nm, 445nm, 488nm, 515nm, 561nm, and 640nm) LUN-F-XL laser engine and a Prime95B camera (Photometrics).

ImageJ was used to quantify the percentage of cells with LRRK2 filaments as previously described. Maximum-intensity projections were generated from *z*-stack confocal images. Using the GFP immunofluorescence signal, transfected cells were identified. Cells were scored for the presence or absence of filaments using both the *z*-projection and *z*-stack micrographs as a guide. To calculate the percentage cells with filaments, the number of cells with filaments was divided by the total number of transfected cells per technical replicate (defined as one 24-well coverslip). Per coverslip, eight fields of view were imaged containing a total of 50 and 150 cells per replicate. The quantification of all cellular experiments come from compiled data collected on at least three separate days. All statistical analyses were performed in GraphPad Prism (9.2; GraphPad Software).

### Western blot analysis and antibodies

For western blot quantification of LRRK2 protein expression and Rab10 phosphorylation, cells were plated on 6-well dishes (200,000 cells per well) 24 h before transfection. Cells were transfected with 500 ng of GFP-LRRK2 construct and 500 ng of GFP-Rab10 using polyethylenimine (PEI, Polysciences). After 36 h, cells were rinsed with ice-cold 1x PBS, pH 7.4 and lysed on ice in RIPA buffer (50 nM Tris pH7.5, 150 mM NaCl, 0.2% Triton X-100, 0.1% SDS, with cOmplete protease inhibitor cocktail and PhoStop phosphatase inhibitor). Lysates were rotated for 15 min at 4°C and clarified by centrifugation at maximum speed in a 4°C microcentrifuge for 15 min. Supernatants were then boiled for 10 minutes in SDS buffer. Experiments were performed in duplicate or triplicate and repeated on at least 3 separate days.

Lysates were run on 4-12% polyacrylamide gels (NuPage, Invitrogen) for 50 minutes at 180V and transferred to polyvinylidene difluoride (Immobilon-FL, EMD Millipore) for 4 h at 200 mA constant current. Blots were rinsed briefly in MilliQ water and dried at room temperature for at least 30 min. Membranes were briefly reactivated with methanol and blocked for 1 h at room temperature in 5% milk (w/v) in TBS. Antibodies were diluted in 1% milk in TBS with 0.1% Tween-20 (TBST). Primary antibodies used for immunoblots were as follows: mouse anti-GFP (Santa Cruz, 1:2500 dilution), rabbit anti-LRRK2 (Abcam, 1:5000 dilution), rabbit anti-GAPDH (Cell Signaling Technology, 1:3000 dilution), and rabbit anti-phospho-T73-RAB10 (Abcam, 1:2500 dilution). Secondary antibodies (1:15000) used for western blots were IRDye goat anti-mouse 680RD and IRDye goat anti-rabbit 780RD (Li-COR). Primary antibodies were incubated overnight at 4°C, and secondary antibodies were incubated at room temperature for 1 h. For quantification, blots were imaged on an Odyssey CLx controlled by Imaging Studio software (v.5.2), and intensity of bands quantified using Image Studio Lite software (v.5.2).

### Cell line

Human 293T cells were obtained from ATCC (CRL-3216) and maintained at 37°C with 5% CO_2_ in Dulbecco’s Modified Eagle Medium (DMEM, Corning) supplemented with 10% fetal bovine serum (FBS, Gibco) and 1% penicillin/streptomycin (PenStrep; Corning). Cells were routinely tested for mycoplasma contamination and were not authenticated after purchase.

### Sequence alignment

Protein sequences of LRRK2 and LRRK1 were obtained from UniProt. Sequence alignments were performed with Clustal Omega web services and annotated using Jalview^75^.

## Extended Data

**Extended Data Fig. 1.**
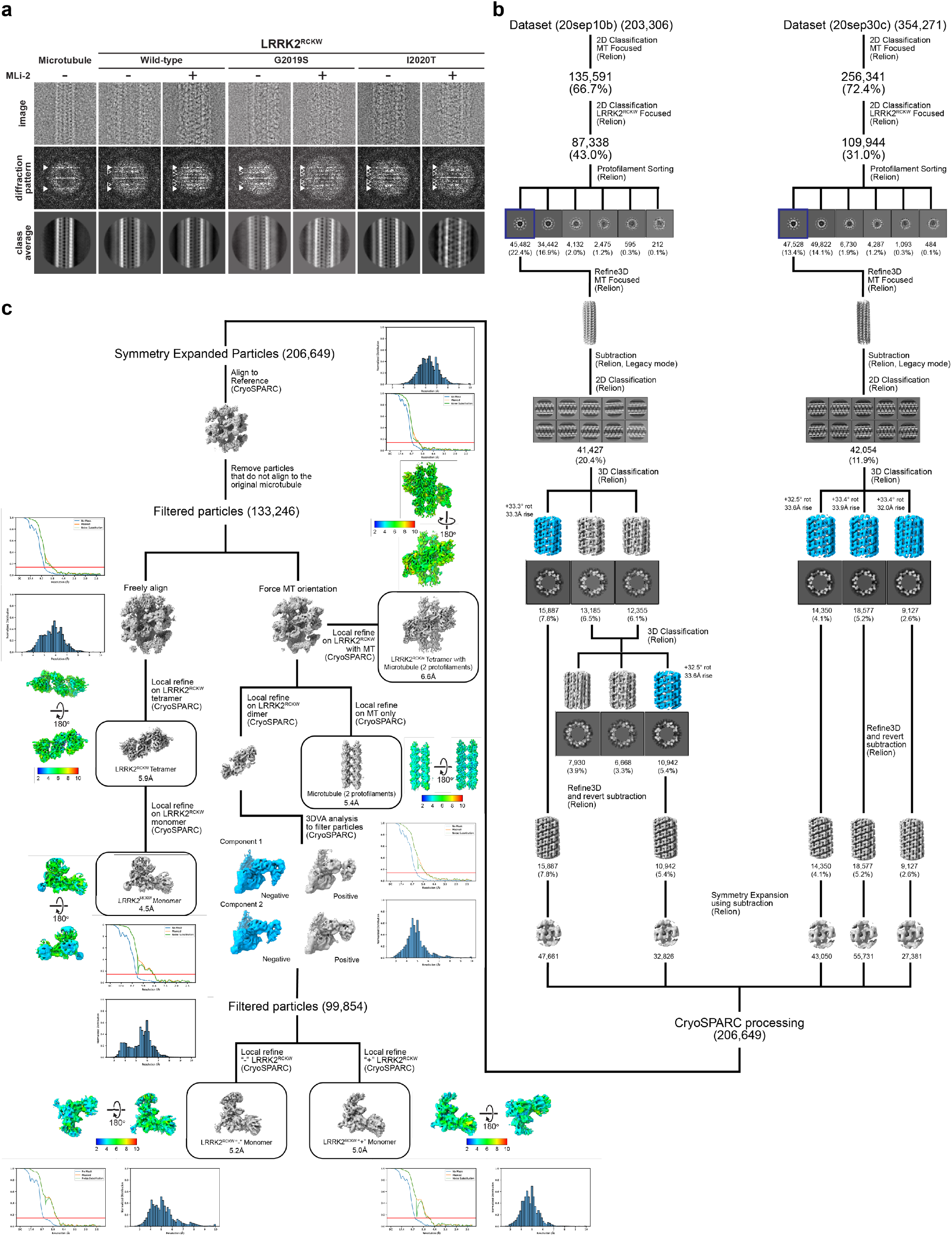
Cryo-EM structure determination of microtubule-associated filaments of LRRK2^RCKW^[I2020T] in the presence of MLi-2. **a**, Optimization of in vitro reconstituted microtubule-associated LRRK2^RCKW^ filaments. Top row, cryo-EM images of an individual microtubule (left) or individual microtubule-associated LRRK2^RCKW^ filaments. Middle, Diffraction patterns calculated from the images above. Arrowheads point to layer lines arising from the microtubule (white) or from the LRRK2^RCKW^ filaments (grey). Bottom, 2D class averages from multiple images equivalent to those shown at the top. The type of LRRK2^RCKW^ (WT, G2019S, or I2020T) and the presence or absence of MLi-2 during filament reconstitution are indicated on top. **b,c**, Schematic of data processing pipeline used to obtain the different reconstructions of the microtubule-associated LRRK2^RCKW^[I2020T] filaments in the presence of MLi-2 (see Methods for details). Local resolution maps, Fourier Shell Correlation plots, and the distribution of voxel resolutions are shown for all reconstructions discussed in the text.

**Extended Data Fig. 2.**
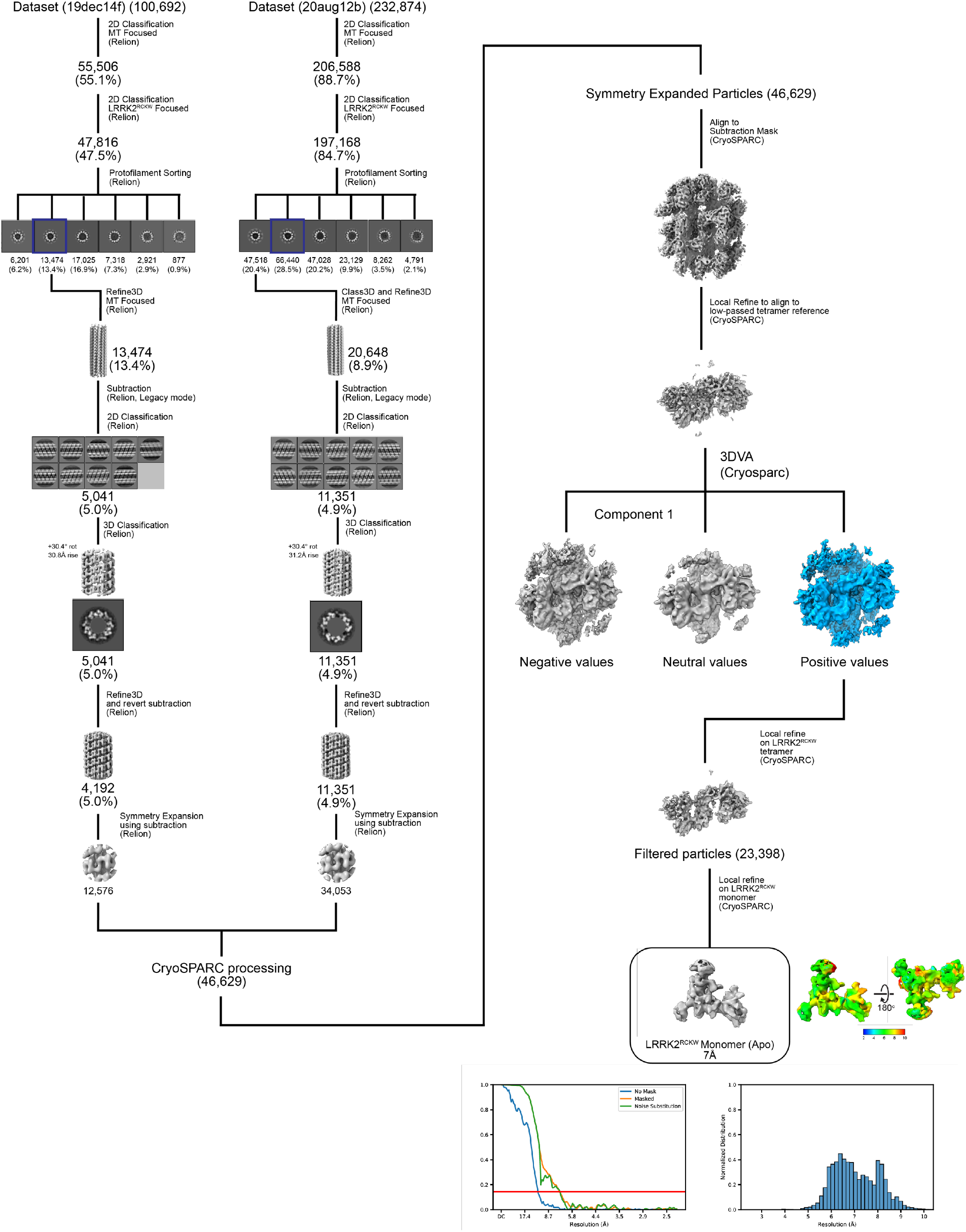
Cryo-EM structure determination of microtubule-associated filaments of LRRK2^RCKW^[I2020T] in the absence of MLi-2. Schematic of data processing pipeline used to obtain the reconstruction of microtubule-associated LRRK2^RCKW^[I2020T] filaments in the absence of MLi-2 (see Methods for details). Local resolution maps, Fourier Shell Correlation plots, and the distribution of voxel resolutions are shown.

**Extended Data Fig. 3.**
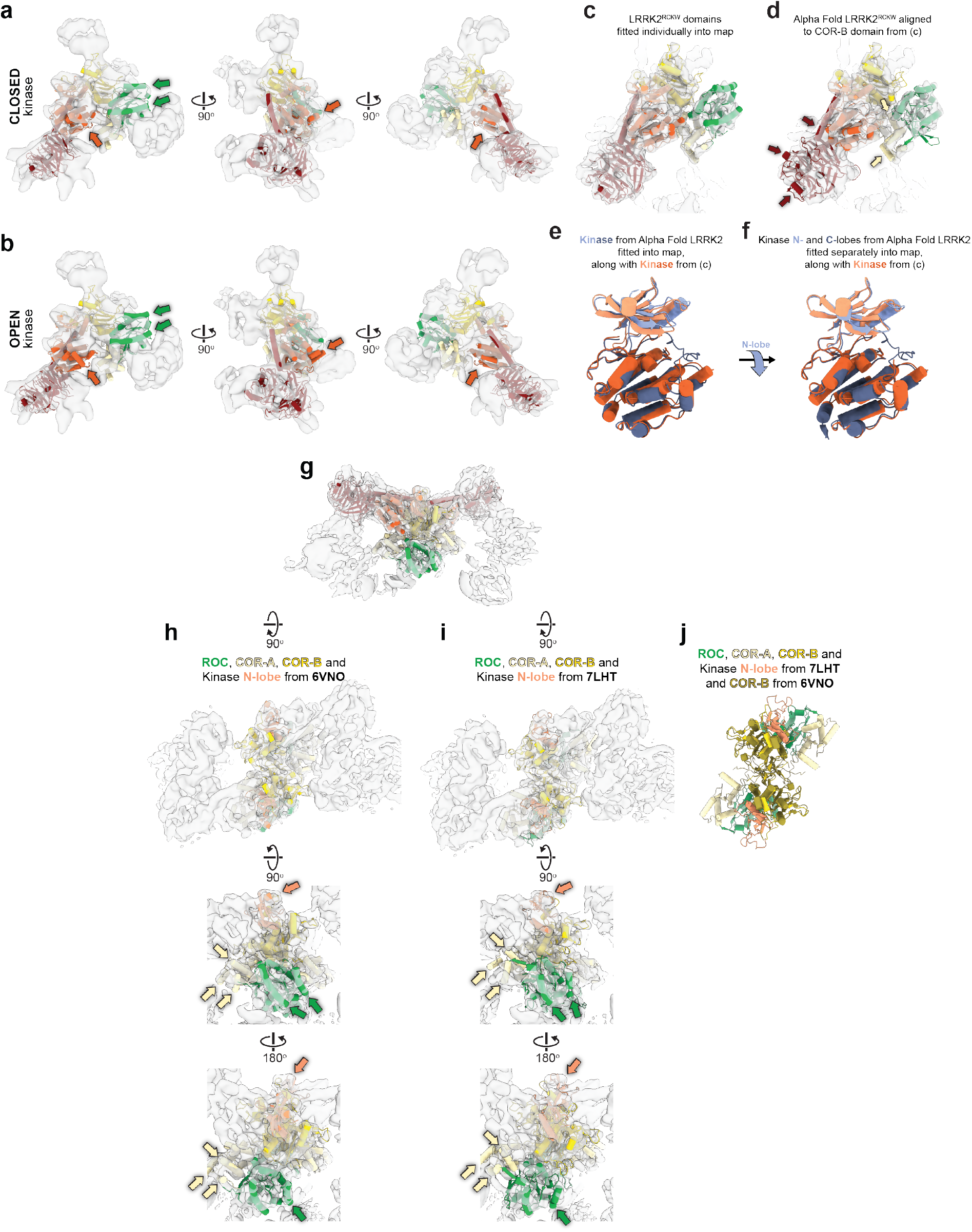
Structural analysis of microtubule-associated filaments of LRRK2^RCKW^[I2020T]. **a**, A model for LRRK2^RCKW^[I2020T] with a closed kinase, obtained by docking the individual domains into the cryo-EM map of LRRK2^RCKW^[I2020T] filaments obtained in the presence of MLi-2, was docked into a cryo-EM map (7Å) of filaments obtained in the absence of the inhibitor (Extended Data Fig. 2). **b**, A model for LRRK2^RCKW^ with an open kinase (PDB:6VNO) was docked into the same map. The colored arrows in (a) and (b) highlight structural elements in the model that protrude from the density when the kinase is in an open conformation. **c**, The LRRK2^RCKW^ domains (ROC, COR-A, COR-B, Kinase N-lobe, Kinase C-lobe, WD40) (PDB:6VNO) were fitted individually into one of the monomers in the cryo-EM map of microtubule-bound LRRK2^RCKW^[I2020T] formed in the presence of MLi-2. **d**, The LRRK2^RCKW^ portion of the AlphaFold model of LRRK2 was aligned to the COR-B domain in (c) and is shown here inside the same cryo-EM map. The colored arrows highlight regions where part of the model protrudes from the density. (Note: there is no arrow pointing to the loop in the ROC domain as this loop was not seen or modeled in the microtubule-bound structure.) **e**, The kinase from the AlphaFold model of LRRK2 was fitted into the cryo-EM map (same as in (d)) and is shown here superimposed on the N- and C-lobes of the kinase as fitted in (c). Note that while the C-lobes superimpose well, the N-lobe fitted individually in (c) is more closed than that modeled in the AlphaFold LRRK2. **f**, The N- and C-lobed of the kinase from the AlphaFold LRRK2 model were now fitted individually into the cryo-EM map (as in (c)), and are shown superimposed on the N- and C-lobes of LRRK2^RCKW^ from (a). The blue arrow between panels (e) and (f) highlights the downward movement of the N-lobe of AlphaFold’s LRRK2 when the two lobes are fitted individually into the cryo-EM map. **g**, The LRRK2^RCKW^ domains (ROC, COR-A, COR-B, Kinase N-lobe, Kinase C-lobe, WD40) (PDB:6VNO) were fitted individually into the central dimer of the cryo-EM map of a tetramer of microtubule-bound LRRK2^RCKW^[I2020T] obtained in the presence of MLi-2. **h,i**, Different closeup views of the map in (g), showing either (h) the ROC, COR-A, COR-B and kinase N-lobe from the LRRK2^RCKW^ model (PDB:6VNO), or (i) the corresponding portion from the structure of full-length LRRK2 (PDB:7LHT) docked as a single body into the cryo-EM map. The colored arrows highlight parts of the model that fit the cryo-EM density better when the domains are fitted in individually (h) rather than as a rigid body (i). **j**, Superposition of the model used in (i) and the COR-B domain from (h) to show that the differences among the ROC, COR-A and N-lobe of the kinase between the two models ((h) and (i)) is not due to major differences at the COR-B:COR-B interface, which is similar.

**Extended Data Fig. 4.**
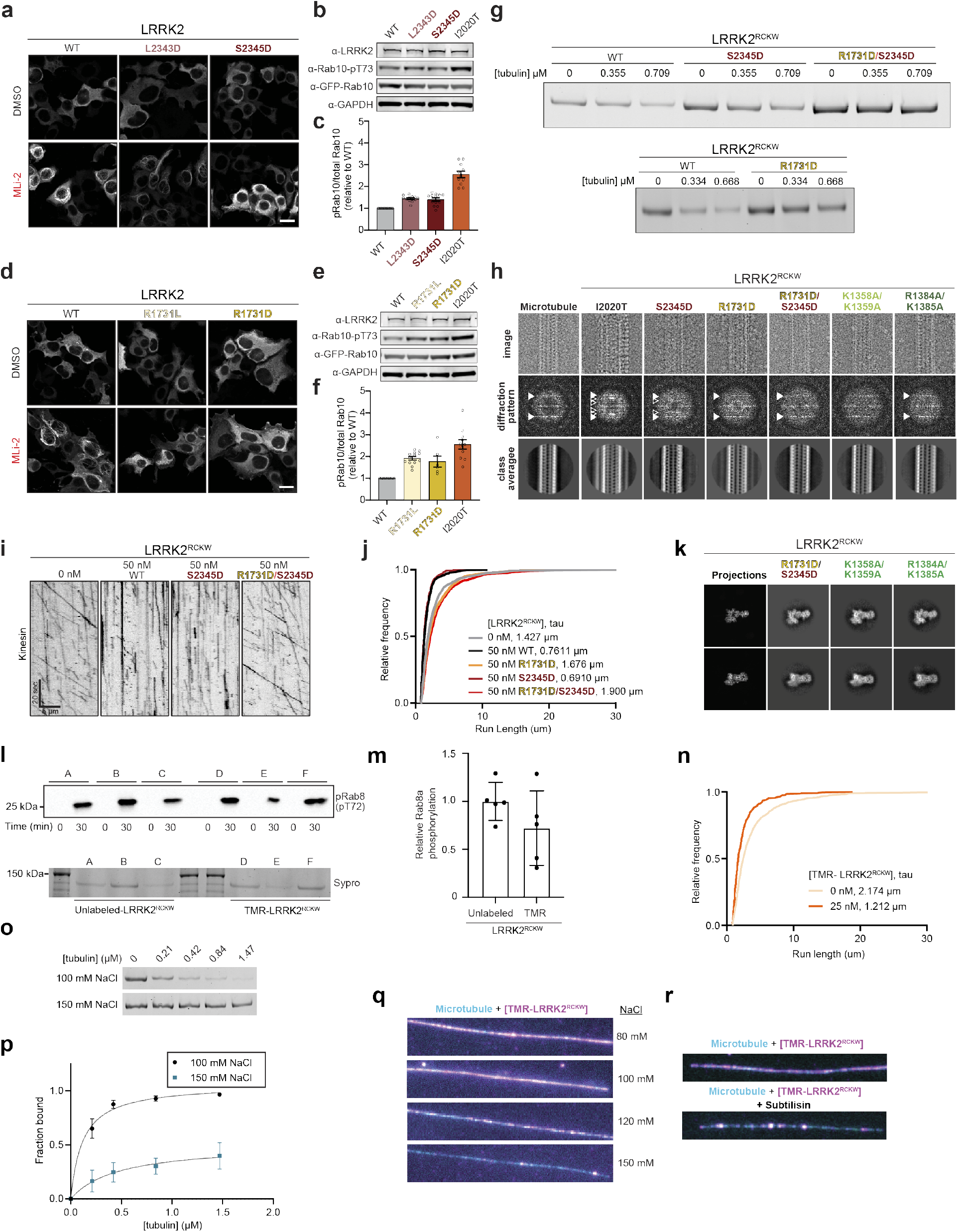
Mechanism of LRRK2^RCKW^ binding to microtubules. **a**, Representative images of 293T cells expressing GFP-LRRK2 (wild type or WD40 mutant as noted on top) and treated with DMSO or 500 nM MLi-2 for 2 hours (as noted at left). Scale bar is 10 μm. **b,c**, Rab10 phosphorylation in 293T cells overexpressing WT LRRK2 or LRRK2 carrying mutations in the WD40 domain. LRRK2[I2020T], which is known to increase Rab10 phosphorylation in cells, was tested as well. 293T cells were transiently co-transfected with the indicated plasmids encoding for GFP-LRRK2 (wild type or mutant) and GFP-Rab10, Thirty-six hours post-transfection the cells were lysed, immunoblotted for phospho-Rab10 (pT73), total GFP-Rab10, and total LRRK2, and developed with LI-COR Odyssey CLx imaging system. Quantification of data in (b) is shown in (c), normalized to wild-type, as mean ± s.e.m. Individual data points represent separate populations of cells obtained across at least three independent experiments. **d**, Representative images of 293T cells expressing GFP-LRRK2 (wild type or COR-B mutant as noted on top) and treated with DMSO or 500 nM MLi-2 for 2 hours (as noted at left). Scale bar is 10 μm. **e,f**, Rab10 phosphorylation in 293T cells overexpressing WT LRRK2 or LRRK2 carrying mutations in the COR-B domain. LRRK2[I2020T] was tested here as well. 293T cells were treated as in (b). Quantification of data in (e) is shown in (f), normalized to wild-type, as mean ± s.e.m. Individual data points represent separate populations of cells obtained across at least three independent experiments. **g**, Representative gel of supernatant from microtubule pelleting assay, used to generate the data shown in Figure 2e. **h**, Cryo-EM analysis of filament formation by LRRK2^RCKW^ mutants. Top row, cryo-EM images of an individual microtubule (left) or combinations of microtubules and LRRK2^RCKW^ mutants. Middle, Diffraction patterns calculated from the images above. Arrowheads point to layer lines arising from the microtubule (white) or from the LRRK2^RCKW^ filaments (grey). Bottom, 2D class averages from multiple images equivalent to those shown at the top. **i**, Example kymographs of single-molecule kinesin motility assays in the presence or absence of 50nM LRRK2^RCKW^ wild-type or indicated mutant. **j**, Cumulative distribution of run lengths for kinesin in the absence or presence of 50 nM LRRK2^RCKW^ (WT or carrying WD40 and/or WD40 and COR-B mutations). The run lengths were not significantly different between 50 nM wild-type and LRRK2^RCKW^ [S2345D], and were significantly between 50 nM wild-type LRRK2^RCKW^ and LRRK2^RCKW^ [R1731D/S2345D] and LRRK2^RCKW^ [R1731D] (Kruskal-Wallis test with Dunn’s post hoc for multiple comparisons). Mean decay constants (tau) are shown. **k**, Comparison of 2D class averages from cryo-EM images of different LRRK2^RCKW^ mutants with the corresponding 2D projection from a LRRK2^RCKW^ molecular model (PDB: 6VNO). Two different views are shown for each mutant. **l**, Representative kinase reaction. Rab8a phosphorylation was measured via western blotting with a phospho-T72-specific Rab8a antibody, and total LRRK2^RCKW^ concentration was measured by Sypro Red staining. Phosphorylation reactions were terminated after 30 minutes. **m**, Quantification of data shown in (l). For each reaction, phospho-Rab8a band intensity (chemiluminescence) was divided by LRRK2^RCKW^ band intensity (Sypro red); for each western blot, an average normalized value was calculated for all replicates of unlabeled LRRK2^RCKW^, and all data was then normalized to this value. **n**, Cumulative distribution of run lengths for kinesin in the absence or presence of 25 nM TMR-LRRK2^RCKW^. The run lengths were significantly different between 0 nM and 25 nM TMR-LRRK2^RCKW^ (Mann-Whitney test). Mean decay constants (tau) are shown. Effect on kinesin motility is similar to previously shown unlabeled LRRK2^RCKW^.**o, p**, Representative microtubule pelleting assay gel for LRRK2^RCKW^ in the presence of 100 mM and 150 mM sodium chloride. Cosedimentation was measured as depletion from supernatant. For each reaction, 200 nM LRRK2^RCKW^ was mixed with a given concentration of microtubules, microtubules were pelleted by high-speed spin, and a gel sample was taken of the supernatant. Quantification of data represented in (o) is shown in (p). Data are mean ± s.d., n=4. The solid line represents a hyperbolic curve fit to the data. **q**, Representative images of coverslip-tethered Alexa Fluor 488-labeled MTs (cyan) bound to 100 nM TMR-LRRK2^RCKW^ (magenta) in the presence of increasing concentrations of sodium chloride, used to generate the data in Figure 3d. **r**, Representative images of untreated (top) and subtilisin-treated (bottom) Alexa Fluor 488-labeled MTs (cyan) bound to 50 nM TMR-LRRK2^RCKW^ (magenta), used to generate the data shown in Figure 3e.

**Extended Data Fig. 5.**
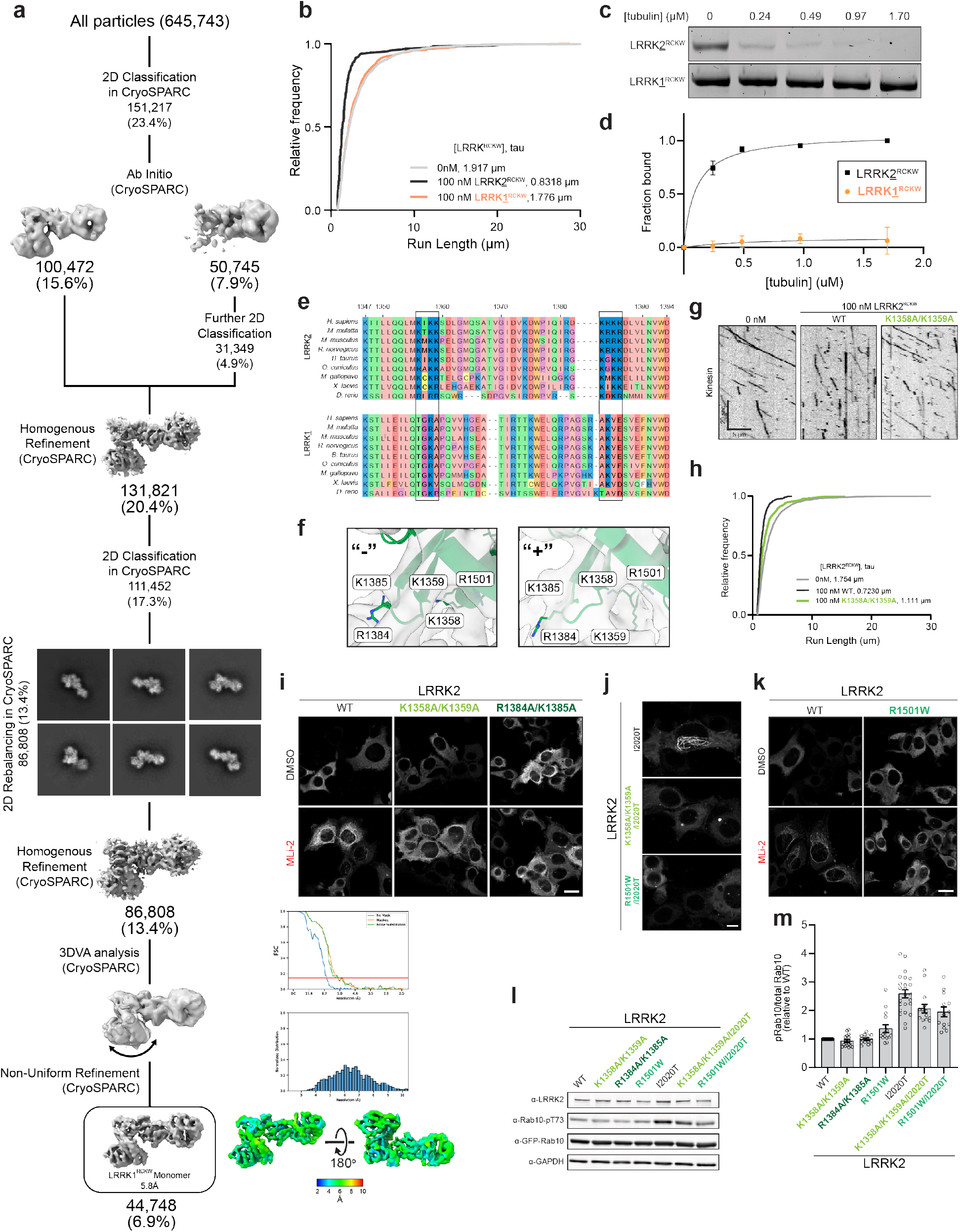
Basic residues within the LRRK2 RoC domain are not conserved in LRRK1 and are involved in LRRK2’s binding to microtubules. **a**, Cryo-EM structure determination of LRRK1^RCKW^. **b**, Cumulative distribution of run lengths for kinesin in the absence or presence of 100 nM LRRK2^RCKW^ or LRRK1^RCKW^. The run lengths were not significantly different between 0 nM and 100 nM LRRK1^RCKW^ conditions (Kruskal-Wallis test with Dunn’s post hoc for multiple comparisons). **c,d**, Representative gel of supernatant from microtubule pelleting assay for 200 nM LRRK2^RCKW^ or LRRK1^RCKW^ with increasing tubulin concentrations. Quantification of data represented in (c) shown in (d). Data are mean ± s.d., n=4. The solid line represents a hyperbolic curve fit to the data. **e**, Sequence alignment of the ROC domains of LRRK2 and LRRK1 across several species made using Clustal Omega. Putative microtubule-contacting residues conserved in LRRK2 but not in LRRK1 are boxed. **f**, Close ups of the basic patches tested in this study, shown in the context of the cryo-EM maps for the and “-” “+” LRRK2^RCKW^ monomers in our reconstruction of the microtubule-associated filaments (Fig. 1e,f). The models shown here correspond to those in Fig. 5c. **g**, Example kymographs of kinesin motility in the presence of 100 nM LRRK2^RCKW^ (wild-type or K1358A/K1359A mutant). **h**, Cumulative distribution of run lengths for kinesin in the absence or presence of 100 nM LRRK2^RCKW^ (WT or carrying ROC mutation). The run lengths were significantly different between 100 nM LRRK2^RCKW^ wild-type and K1358A/K1359A mutant (Kruskal-Wallis test with Dunn’s post hoc for multiple comparisons). Mean decay constants (tau) are shown. **i**, Representative images of 293T cells expressing GFP-LRRK2 (wild type or ROC mutant as noted on top) and treated with DMSO or 500 nM MLi-2 for 2 hours (as noted at left), corresponding to data plotted in Fig. 5f. Scale bar is 10 μm. **j**, Representative images of 293T cells expressing GFP-LRRK2 (I2020T, I2020T/ROC mutant, or I2020T/R1501W, as noted at left), corresponding to data plotted in Fig. 5g,i. Scale bar is 10 μm. **k**, Representative images of 293T cells expressing GFP-LRRK2 (wild type or R1501W mutant as noted on top) and treated with DMSO or 500 nM MLi-2 for 2 hours (as noted at left), corresponding to data plotted in Fig. 5h. Scale bar is 10 μm. **l,m**, Rab10 phosphorylation in 293T cells overexpressing WT LRRK2 or LRRK2 carrying indicated mutations in the ROC domain. LRRK2[I2020T], which is known to increase Rab10 phosphorylation in cells, was tested as well. 293T cells were transiently transfected with the indicated plasmids encoding for GFP-LRRK2 (wild type or mutant) and GFP-Rab1O, Thirty-six hours post-transfection the cells were lysed, immunoblotted for phospho-Rab10 (pT73), total GFP-Rab10, and total LRRK2, and developed with LI-COR Odyssey CLx imaging system. Quantification of data in (l) is shown in (m), normalized to wild-type, as mean ± s.e.m. Individual data points represent separate populations of cells obtained across at least three independent experiments.

**Extended Data Fig. 6.**
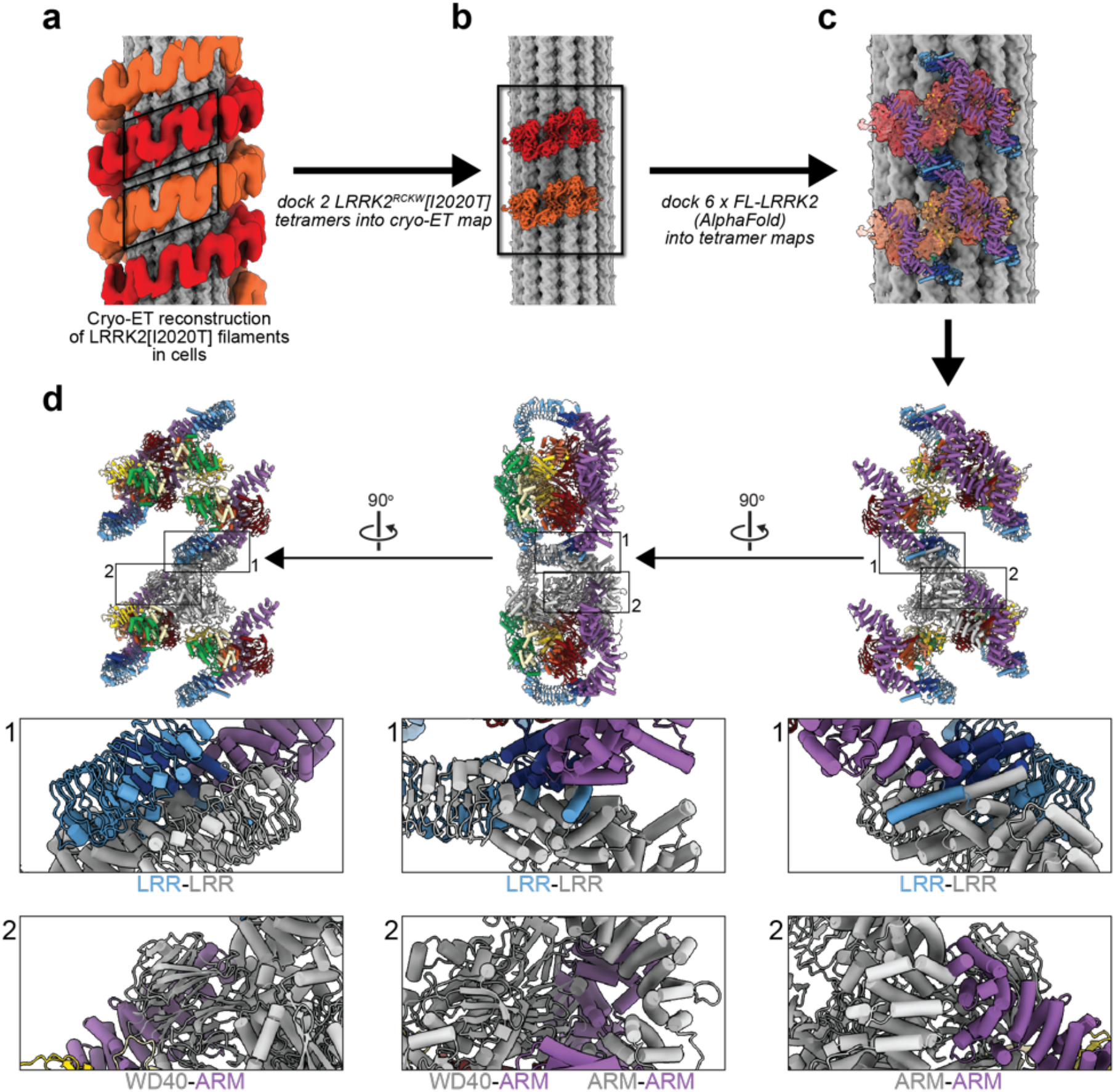
Modeling of full-length LRRK2 into the cryo-ET reconstruction of microtubule-associated LRRK2[I2020T] filaments in cells. **a**, Cryo-ET reconstruction of microtubule-associated LRRK2[I2020T] filaments in cells (Villa). The LRRK2 strands that form the double-helical filaments are shown in light and dark orange. For this figure, the density corresponding to the microtubule was replaced with a 10Å representation of a molecular model of a microtubule. **b**, We docked copies of our 5.9Å reconstruction of a LRRK2^RCKW^[I2020T] tetramer from the microtubule-associated filaments (Fig. 1b,c) into the regions indicated by the parallelograms in (a). **c**, Next, we docked two copies of the AlphaFold model of full-length LRRK2 (AF-Q5S007), which is in the active state, as is the case with LRRK2^RCKW^[I2020T] in our filaments, into each of the 5.9Å maps. The pairs of AlphaFold models in each map correspond to the COR-B:COR-B dimer. This panel shows a region corresponding to the rectangle in (b). **d**, Three different views of the models docked in (c). Below each model, close-ups show regions where adjacent filaments clash. These clashes involve a domain in the N-terminal repeats of one LRRK2, and either the same domain on another LRRK2, or the WD40 domain. For clarity, one of the LRRK2’s is shown in grey instead of the standard rainbow coloring.

**Supplementary Table 1.**

|  | **LRRK2<sup>RCKW</sup> + MT**<br>**+MLi-2 (Helical)** | **LRRK2<sup>RCKW</sup> + MT**<br>**+MLi-2 (Tetramer**<br>**only)** | **LRRK2<sup>RCKW</sup> + MT**<br>**+MLi-2 (Tetramer +**<br>**MT)** |
| --- | --- | --- | --- |
|  | EMD-25649<br>EMPIAR-10925 | EMD-25664<br>EMPIAR-10925 | EMD-25658<br>EMPIAR-10925 |
| **Data Collection** |  |  |  |
| Microscope | Talos Arctica | Talos Arctica | Talos Arctica |
| Camera | K2 summit | K2 Summit | K2 Summit |
| Magnification | 36,000 | 36,000 | 36,000 |
| Voltage (kV) | 200 | 200 | 200 |
| Total electron exposure (e-/Å<sup>2</sup>) | 55 | 55 | 55 |
| Defocus Range (um) | 0.5-2.5 | 0.5-2.5 | 0.5-2.5 |
| Pixel Size (Å/pixel) | 1.16 | 1.16 | 1.16 |
| Number of dataset (no.) | 2 | 2 | 2 |
| **Reconstruction** |  |  |  |
| Total Extracted picks (no.) | 354,271 | 206,649 (symmetry<br>expanded) | 206,649 (symmetry<br>expanded) |
| Final Particles (no.) | 14,350 | 133,246 | 133,246 |
| Symmetry | +32.5° rot, 33.3Å<br>rise | C1 | C1 |
| Resolution (global) (Å) |  |  |  |
| FSC 0.143 | 18 | 5.9 | 6.6 |
| Local resolution range (Å) | N/A | 3.5-9 | 3.7-9.5 |
| Map sharpening *B*-factor | N/A | -339 | -326 |
|  | **LRRK2<sup>RCKW</sup> + MT**<br>**+MLi-2**<br>**(Microtubule only)** | **LRRK2<sup>RCKW</sup> + MT**<br>**Apo (Monomer**<br>**only)** | **LRRK1<sup>RCKW</sup>** |
|  | EMD-25908<br>EMPIAR-10925 | EMD-25674<br>EMPIAR-10924 | EMD-25672<br>EMPIAR-10921 |
| **Data Collection** |  |  |  |
| Microscope | Talos Arctica | Talos Arctica | Talos Arctica |
| Camera | K2 summit | K2 Summit | K2 Summit |
| Magnification | 36,000 | 36,000 | 36,000 |
| Voltage (kV) | 200 | 200 | 200 |
| Total electron exposure (e-/Å<sup>2</sup>) | 55 | 50-55 | 55 |
| Defocus Range (um) | 0.5-2.5 | 0.5-2.5 | 1.2-1.8 |
| Pixel Size (Å/pixel) | 1.16 | 1.16 | 1.16 |
| Number of dataset (no.) | 2 | 2 | 4 |
| **Reconstruction** |  |  |  |
| Total Extracted picks (no.) | 206,649 (symmetry<br>expanded) | 46,629 (symmetry<br>expanded) | 645,743 |
| Final Particles (no.) | 133,246 | 23,398 | 44,748 |
| Symmetry | C1 | C1 | C1 |
| Resolution (global) (Å) |  |  |  |
| FSC 0.143 | 5.4 | 7.0 | 5.8 |
| Local resolution range (Å) | 2.6-9.0 | 4.8-10.0 | 3.5-10 |
| Map sharpening *B*-factor | -235 | -293 | -327 |

|  | **LRRK2<sup>RCKW</sup> + MT**<br>**+MLi-2 (Focused**<br>**on Kinase)** | **LRRK2<sup>RCKW</sup> + MT**<br>**+MLi-2 (Minus end)** | **LRRK2<sup>RCKW</sup> + MT**<br>**+MLi-2 (Plus end)** |
| --- | --- | --- | --- |
|  | EMD-25897<br>EMPIAR-10925 | 7THY<br>EMD-25906<br>EMPIAR-10925 | 7THZ<br>EMD-25907<br>EMPIAR-10925 |
| **Data Collection** |  |  |  |
| Microscope | Talos Arctica | Talos Arctica | Talos Arctica |
| Camera | K2 summit | K2 Summit | K2 Summit |
| Magnification | 36,000 | 36,000 | 36,000 |
| Voltage (kV) | 200 | 200 | 200 |
| Total electron exposure (e-/Å<sup>2</sup>) | 55 | 55 | 55 |
| Defocus Range (um) | 0.5-2.5 | 0.5-2.5 | 0.5-2.5 |
| Pixel Size (Å/pixel) | 1.16 | 1.16 | 1.16 |
| Number of dataset (no.) | 2 | 2 | 2 |
| **Reconstruction** |  |  |  |
| Total Extracted picks (no.) | 206,649 (symmetry<br>expanded) | 206,649 (symmetry<br>expanded) | 206,649 (symmetry<br>expanded) |
| Final Particles (no.) | 133,246 | 99,854 | 99,854 |
| Symmetry | C1 | C1 | C1 |
| Resolution (global) (Å) |  |  |  |
| FSC 0.143 | 4.5 | 5.2 | 5.0 |
| Local resolution range (Å) | 3.0-8.0 | 2.6-9.0 | 2.9-7.0 |
| Map sharpening B-factor | -146 | -200 | -200 |
| **Refinement** |  |  |  |
| Initial model used | N/A | Q5S007 (AlphaFold) | Q5S007 (AlphaFold) |
| Model resolution (Å) | N/A | 5.4 (average) | 5.3 (average) |
| Resolution method | N/A | Q-score | Q-score |
| Model resolution range (Å) | N/A | 3.0-8.6 | 3.5-8.0 |
| Model refinement package | N/A | Rosetta, Isolde | Rosetta, Isolde |
| Number of models | N/A | 5 | 5 |
| Nonhydrogen atoms | N/A | 1584 (per model) | 1584 (per model) |
| Protein residues | N/A | 194 (per model) | 194 (per model) |
| B factors (Å<sup>2</sup>) |  |  |  |
| Protein residues | N/A | -219 (average) | -216 (average) |
| R.m.s. deviations |  |  |  |
| Bond Lengths (Å) | N/A | 0.019 (average) | 0.020 (average) |
| Bond angles (°) | N/A | 1.907 (average) | 2.023 (average) |
| **Validation** |  |  |  |
| MolProbity score | N/A | 1.49 (average) | 1.54 (average) |
| Clashscore | N/A | 3.52 (average) | 3.96 (average) |
| Poor rotamers (%) | N/A | 0 (average) | 0 (average) |
| Ramachandran plot |  |  |  |
| Favored (%) | N/A | 94.5 (average) | 94.6 (average) |
| Allowed (%) | N/A | 4.9 (average) | 4.8 (average) |
| Disallowed (%) | N/A | 0.6 (average) | 0.6 (average) |
| Map CC (CCmask) | N/A | 0.63 (average) | 0.61 (average) |

**Supplementary Table 2:**
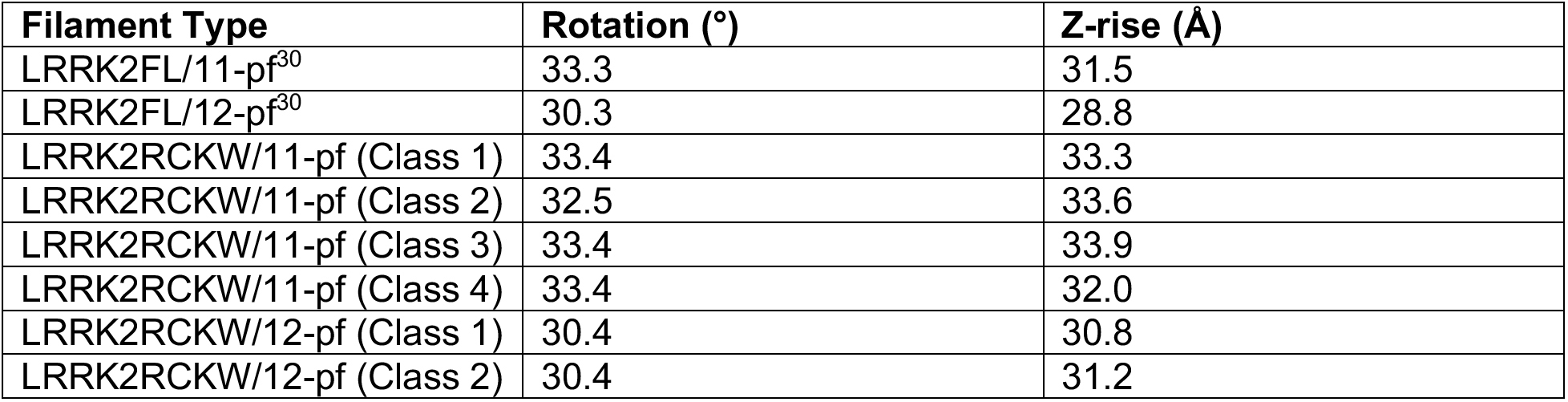
Parameters observed for LRRK2^RCKW^ filaments compared to published LRRK2FL filaments.

